# Enriched Single-Nucleus RNA-Sequencing reveals unique attributes of distal convoluted tubule cells

**DOI:** 10.1101/2023.09.05.555810

**Authors:** Xiao-Tong Su, Jeremiah V. Reyes, Anne E. Lackey, Yujiro Maeoka, Ryan J. Cornelius, James A. McCormick, Chao-Ling Yang, Hyun Jun Jung, Paul A. Welling, Jonathan W. Nelson, David H. Ellison

**Affiliations:** Division of Hypertension and Nephrology, School of Medicine, Oregon Health and Science University, Oregon, USA; Department of Biochemistry and Molecular Biology, College of Medicine, University of the Philippines Manila, Manila, Philippines; Division of Nephrology, Department of Medicine, Johns Hopkins University School of Medicine, Baltimore, USA; Department of Physiology, Johns Hopkins University School of Medicine, Baltimore, USA; Oregon Clinical and Translational Research Institute, Oregon Health and Science University, Oregon, USA; Renal Section, VA Portland Healthcare System, Oregon, USA

## Abstract

**Background:** The distal convoluted tubule (DCT) comprises two subsegments, DCT1 and DCT2, with different functional and molecular characteristics. The functional and molecular distinction between these segments, however, has been controversial.

**Methods:** To understand the heterogeneity within the DCT population with better clarity, we enriched for DCT nuclei by using a mouse line combining “Isolation of Nuclei TAgged in specific Cell Types” and NCC (sodium chloride cotransporter)-driven inducible Cre recombinase. We sorted the fluorescently labeled DCT nuclei using Fluorescence-Activated Nucleus Sorting, and performed single nucleus transcriptomics.

**Results:** Among 25,183 DCT cells, 70% were from DCT1 and 30% from DCT2. Additionally, there was a small population (<1%) enriched in proliferation-related genes, such as*Top2a, Cenpp, and Mki67.* Both DCT1 and DCT2 express NCC, magnesium transport genes are more abundant along DCT1; whereas calcium, electrogenic sodium and potassium transport genes are more abundant along DCT2. The transition between these two segments are gradual with a transitional zone where DCT1 and DCT2 cells are interspersed. The expression of the homeobox genes is not consistent between all DCT cells, suggesting that they develop along different trajectories.

**Conclusion:** Transcriptomics analysis of an enriched rare cell population using genetically targeted approach offers better clarification of the function and classification. The DCT segment is short, yet, can be separated into two sub-types that serve distinct functions, and are speculated to derive from different origins during development.

**Significance Statement:** High-resolution snRNAseq data indicate a clear separation between primary sites of calcium and magnesium handling within DCT. Both DCT1 and DCT2 express *Slc12a3*, but these subsegments serve distinctive functions, with more abundant magnesium handling genes along DCT1 and more calcium handling genes along DCT2. The data also provides insight into the plasticity of the distal nephron-collecting duct junction, formed from cells of separate embryonic origins. By focusing/changing gradients of gene expression, the DCT can morph into different physiological cell states on demand.

## Introduction

The mammalian kidneys play a critical role in regulating of electrolyte balance and maintaining blood pressure. At least 14 kidney tubule segments have been described within the nephron, of which the distal convoluted tubule (DCT) is the shortest; yet this short segment plays a crucial role in various homeostatic processes, including sodium, potassium, calcium and magnesium handling. The DCT comprises two subsegments commonly called DCT1 and DCT2, with different functional and molecular characteristics ^1–3^. The functional and molecular distinction between these segments, however, has been controversial ^4^ and inconsistently delineated within single-cell transcriptomic datasets from kidney tissue ^5–8^. This likely reflects the fact that DCT cells comprise only a small percentage of kidney cortical tissue. The best study to date, therefore, enriched for DCT cells using cell-surface markers by Knepper and colleagues, thus clearly identifying DCT1 and DCT2 cells ^9^; in that study, however, the proportions of DCT1 and DCT2 cells were not similar to proportions reported using more standard approaches^10, 11^.

They also identified a cluster of DCT-derived cells with features consistent with a proliferative phenotype ^10^. The ‘distal tubule’, including the DCT, connecting tubule (CNT), and cortical collecting duct (CCD) is embryologically unique, as it develops from fusion of the nephrogenic blastema with the ureteric bud. Fate mapping experiments suggest that cells from a distal nephron lineage reprogram during development to resemble cells of ureteric lineage, through a process of mutual induction ^12^. This origin may explain the difficulty, despite decades of work, in establishing a single site at which nephron and bud join, but it also has been suggested as a reason that this segment is uniquely plastic, in response to physiological perturbations. To date, the failure to recapitulate fusion of nephron and bud has been a limitation towards kidney organoid development ^13^.

As the DCT comprises only a small percentage of renal cortical cells, the information gained from single cell analysis of whole kidney tissue is often limited. Prior efforts to characterize DCT cells have been limited by the rarity of these cells within the kidney. To address this, Chen *et al.* enriched for DCT cells using a cell surface marker-based fluorescence-activated cell sorting protocol and analyzed the enriched distal nephron population using the single cell RNA sequencing (scRNA-Seq); they not only confirmed the existence of DCT1 and DCT2, but also found a population of cells that were enriched in cell cycle-and cell proliferation-associated genes ^9^. Of note, however, the proportion of DCT1 to DCT2 was 13:1 in their study, which differs from the ratio observed in morphological studies^11^. Thus, we utilized a highly enriched population of DCT cells generated using the Isolation of Nuclei TAgged in specific Cell Types (INTACT) system for single-nucleus RNA-sequencing (snRNA-Seq) to understand the heterogeneity within the DCT population with better clarity.

## Methods

### Study approval

Animal studies were approved by the Oregon Health and Science University IACUC (protocol IP00286).

### Mouse models

All mice were housed under standard conditions, under light/dark cycle (12:12hr), with free access to food and drinking water. NCC-Cre-INTACT mice were used for snRNA-Seq experiments. The Cre/LoxP technology was used to label DCT nuclei with a nuclear GFP, which allowed for subsequent FANS enrichment. This system, called INTACT (Isolation of Nuclei TAgged in specific Cell Types), fluorescently labels the nuclei from genetically targetable populations of cells. This is achieved by tagging the C-terminus of SUN1, a nuclear membrane protein, with two tandem copies of superfolder GFP and six copies of the Myc epitope (SUN1-sfGFP-Myc)^14, 15^. The INTACT mice were crossed with the NCC-CreERT2 mice ^16^ to create a mouse line expressing sfGFP at the inner nuclear membrane in DCT cells upon tamoxifen induction. Three female NCC-Cre-INTACT mice were i.p. injected with 1 mg daily for five days followed by a ten-day induction period to induce the sfGFP expression. Water was provided *ad libitum*.

### Nuclei isolation from mouse kidney cortex and FANS

Kidneys were removed after PBS washout via aortic perfusion, and the cortex was dissected and then snap-frozen in liquid nitrogen. Frozen tissue was stored at −80 °C until subsequent tissue processing. We modified from the TST (INNER cell) nuclei extraction ^17^ and the kidney nuclei isolation protocol ^5, 18^. The nuclei isolation buffer (NIB) contains 146 mM NaCl, 10 mM Tris-HCl (pH 7.5), 1 mM CaCl_2_, 21 mM MgCl_2_, 0.03% Tween-20, 0.01% BSA, and 1 tablet of cOmplete ULTRA per 10 ml NIB. The samples were ground for 30 times in 2ml NIB1 (4mL NIB + 20µl RNasin Plus + 20µl SUPERaseIN) with a 2ml Dounce grinder and a loose pestle, then homogenate was passed through a 200 µm strainer. The homogenate was ground for 15 times with a tight pestle, then 2 ml NIB1 was added and the homogenate was incubated for 5 min on ice. The homogenate was passed through a 40 µm strainer, then centrifuged at 500g for 5 min at 4°C. The pellet was resuspended in 4 ml NIB2 (4mL NIB + 4µl RNasin Plus + 4µl SUPERaseIN) and the suspension was incubated on ice for 5 min, then centrifuged at 500g for 5 min at 4°C. The pellet was resuspended in 1.5 ml of nuclei resuspension buffer (NSB, 10mL DPBS + 10µl RNasin Plus), then centrifuged at 500g for 5 min at 4°C. Repeat this step, then the suspension was passed through a 5 µm strainer. The suspension was centrifuged at 500g for 5 min at 4°C, and the pellet was resuspended in 10 ml NSB. The nuclei suspension was mixed with 5µl Vybrant Ruby stain and incubated on ice for 15 min. The nuclei were sorted in 500 µl final resuspension buffer (FSB, 1ml DPBS with 1% BSA + 5 µl Protector RNase inhibitor) with a low flow rate and pressure to ensure high viability. Two main gates were used. A 561+683 emission for the ruby stain and a 488 530/40 emission for GFP. Low trigger pulse width was used as singlet discriminator. One hundred thousand nuclei were collected and centrifuged at 500g for 5 min at 4°C. The top supernatant was carefully removed, and 50 µl volume was left to resuspend the nuclei pellet. Ten microliters of nuclei were mixed with 10 µl Trypan Blue and then loaded on the Fuchs-Rosenthal disposable hemocytometer for counting. Nuclei at 700-1,200 nuclei/µl were directly loaded to 10X chips for Gel Bead-In Emulsions (GEM) generation.

### SnRNA-Seq

The snRNA-Seq was performed using a Chromium Next GEM Single Cell 3 Reagent Kit v3.1 (10x Genomics). Single nuclei were partitioned in droplets with single Gel Beads, which contained primers with cell-tagging indexes. Single nucleus suspensions with concentration of at least 300/µl were loaded targeting 10,000 nuclei per sample. The resulting cDNA was profiled on a Bioanalyzer NanoChip (Agilent), then used as a template for library preparation according to the reagent kit protocol. The final libraries were profiled on a Tapestation D1000 tape (Agilent) and quantified using real time PCR (Kapa Biosystems) on a StepOnePlus Real Time PCR workstation (Thermo/ABI). Libraries were sequenced on a NovaSeq 6000 (Illumina). FASTQ files were prepared using bcl2fastq (Illumina) and then aligned to a reference genome using Cellranger v. 6.1.2 (10x Genomics). Reads were mapped to both exonic and intronic regions to include the pre-mRNA transcriptome. Library preparation and sequencing were done by the OHSU Integrated Genomics Laboratory, RRID:SCR_022651.

### SnRNA-Seq Data Analysis

#### Ambient RNA, doublet, and mitochondrial feature removal

Ambient RNA contamination was estimated and removed using SoupX v. 1.6.1 ^19^. Cells with cleaned-up reads were then subjected to doublet removal using DoubletFinder ^20^. Mitochondrial features were manually removed by excluding all features starting with *mt-*^21^. These processing steps were done separately for each sample. Count matrices now only containing singlets were loaded to Seurat v. 4.0 for merging and further filtering ^22^. Nuclei with the features > 4,000 or < 500, and counts > 10,000 were considered low quality nuclei and were filtered out. The filtered Seurat object was then inputted to SCTransform for normalization and variance stabilization ^23^. The normalized count matrix was integrated using Seurat’s built-in *FindIntegrationAnchors*.

#### Batch effect correction, and cell clustering

Dimensional reduction analysis was performed using Principal Component Analysis (PCA). Clusters were generated using the built-in *FindNeighbors* and *FindClusters* Seurat commands employing the first 50 PCA dimensions with a 0.15 clustering resolution. The Uniform Manifold Approximation and Projection (UMAP) technique was used for 2D projection and visualization. Clusters were annotated based on selected canonical gene markers and the top 5 genes in the cluster as defined using *FindAllMarkers*. DCT clusters, namely DCT1 (early distal convoluted tubule), DCT2 (late distal convoluted tubule), and Prolif (proliferating cell population), were manually separated and re-integrated for downstream analyses. The final Seurat object from three animals had a total of 21280 features from 25183 nuclei.

#### SnRNA-seq differential gene analysis

Differential gene expression analysis was done with *FindMarkers* using the normalized and corrected values stored in the sct slot employing the DESeq2 mode ^24^. The final differentially expressed gene list was filtered based on minimum detection rate (min.pct) at 10%, average fold difference of 1.5 (log fold difference > 0.3219), and adjusted p-value of 0.05.

#### Cellular Identity and Functional Scores

Using Seurat’s *AddModuleScore* function, we computed scores using the average expression levels of a set of genes on a single cell level. *DCT1 score* was computed from the average expression of the top 10 genes (*Erbb4*, *Egf*, *Trpm7*, *Fgf13*, *Col5a2*, *Umod*, *Ptgfr*, *Stk32b*, *Rtl4*, *Abca13*) distinguishing DCT1 from DCT2 based after running *FindMarkers* in our control dataset. The same was done to compute for the DCT2 score using the following gene set – *Slc8a1*, *Arl15*, *Calb1*, *Slc2a9*, *Phactr1*, *Gls*, *S100g*, *Kl*, *Klk1*, and *Egfem1. Mg score* was computed from the average expression of a curated list of known magnesiotropic genes expressed in the DCT that have reported magnesium derangement phenotype namely, *Trpm6*, *Trpm7*, *Egf*, *Umod*, *Prox1*, *Fxyd2*, *Cnnm2*, *Slc41a3* ^25–33^. *Ca score* was computed from the average expression of a curated list of known calciotropic genes expressed in the DCT that have reported calcium derangement phenotype namely, *Trpv5*, *Calb1*, *Slc8a1*, *S100g*, *Vdr*, and *Ryr2* ^34–40^.

#### Data Visualization

Dimension, feature, dot, and violin plots were generated from built in Seurat visualization functions. Score density plots were generated using the density function of ggplot2 ^41^. Kernel density estimation was done to fit a smooth curve reflecting the histogram of the module score distribution.

#### Pseudotime analysis

The Seurat object was converted to a Monocle cell data set using *SeuratWrappers* ^42–45^. Pseudotime was computed using *monocle3* and was used to plot nuclei into a normalized expression heatmap.

#### Pathway Analysis

The differentially expressed gene list was subjected to overrepresentation analysis using the *enrichGo* function of *ClusterProfiler* ^46^. Gene symbols were converted to ENTREZIDs using the Biological ID translator feature. Upregulated pathways were defined as pathways corresponding to the genes with actual log fold difference of > 1.25 and downregulated pathways with values < −1.25.

### Immunofluorescence

Mice were anesthetized with a ketamine-xylazine-acepromazine cocktail (50:5:0.5 mg/kg). The right kidney was tied off, removed, and flash frozen in liquid nitrogen for isolation of protein. The left kidney was perfusion fixed with retrograde abdominal aortic perfusion of 3% paraformaldehyde in PBS (pH 7.4). After perfusion, the left kidney was removed, dissected, and cryopreserved in 800 mOsm sucrose in PBS overnight before being embedded in Tissue-Tek OCT compound (Sakura Finetek, Torrance, CA). Slides were prepared by cutting 5-µm sections and were stored at −80°C until use. Immunofluorescent staining was prepared as follows. Slides were incubated with 0.5% Triton X-100 in PBS for 30 min. Sections were then blocked with 5% milk in PBS for 30 min followed by incubation with primary antibody, diluted in blocking buffer, for 1 h at room temperature or overnight at 4°C. Sections were washed with PBS three times and incubated with fluorescent dye-conjugated secondary antibody, diluted in blocking buffer, for 1 h at room temperature. Sections were washed with PBS three times and stained with DAPI before being mounted with ProLong® Diamond Antifade Mountant (Thermo Fisher Scientific, Carlsbad, CA). For micro-dissected tubule immunofluorescence, kidneys were perfused and digested in L-15 medium with 1 unit/ml type II collagenase at 37 °C for 30 minutes. The micro-dissected tubules were transferred to 5 mm x 5 mm cover slip coated with poly-D-lysine. The tubules were fixed with formalin for 30 minutes on ice and stained following the same protocol described above. Images were captured using a KEYENCE BZ-X800 microscope (Itasca, IL). Image processing was completed using FIJI software (National Institutes of Health, Bethesda, MD).

### Antibodies and chemicals

A list of antibodies and chemicals including vendor and catalog numbers is provided in Supplemental table 1.

### Data availability

Sequencing data are deposited in GEO under accession number GSE228367. To ensure data accessibility to non-bioinformaticians, we made the DCT snRNA-Seq data available for further exploration via an interactive web tool generated using ShinyCell ^47^ at https://ellisonlab.shinyapps.io/dct_shinycell/.

### Code availability

All analysis code is available on GitHub at: https://github.com/OHSU-NHT/Su_dct_2023.

## Results

### 1) Enriched snRNA-Seq confirms segmentation of the distal convoluted tubule

To determine the heterogeneity of DCT cells, we took advantage of single-cell transcriptomics and performed snRNA-Seq. To enable more robust enrichment of nuclei from DCT cells and introduce less processing artifacts, thereby increasing the cytological resolution, we bred a DCT-specific inducible NCC-Cre recombinase mouse line ^16^ with the INTACT^flx/flx^ ^15^ mouse line to generate DCT-INTACT mice. These mice express a green fluorescent protein conjugated to the nuclear envelope protein Sun1 in cells that also express NCC (**Figure 1A**). We then isolated nuclei from flash-frozen kidneys of these mice by combining approaches from Humphreys and Regev ^5, 17, 18^, and then enriching for DCT nuclei using fluorescence-activated nuclei sorting (FANS) of the signal from GFP and DNA (**Figure 1B, S1**). We confirmed the integrity of the nuclei following FANS (**Figure 1C**) prior to processing through the 10x chromium controller for sequencing.

**Figure 1.**
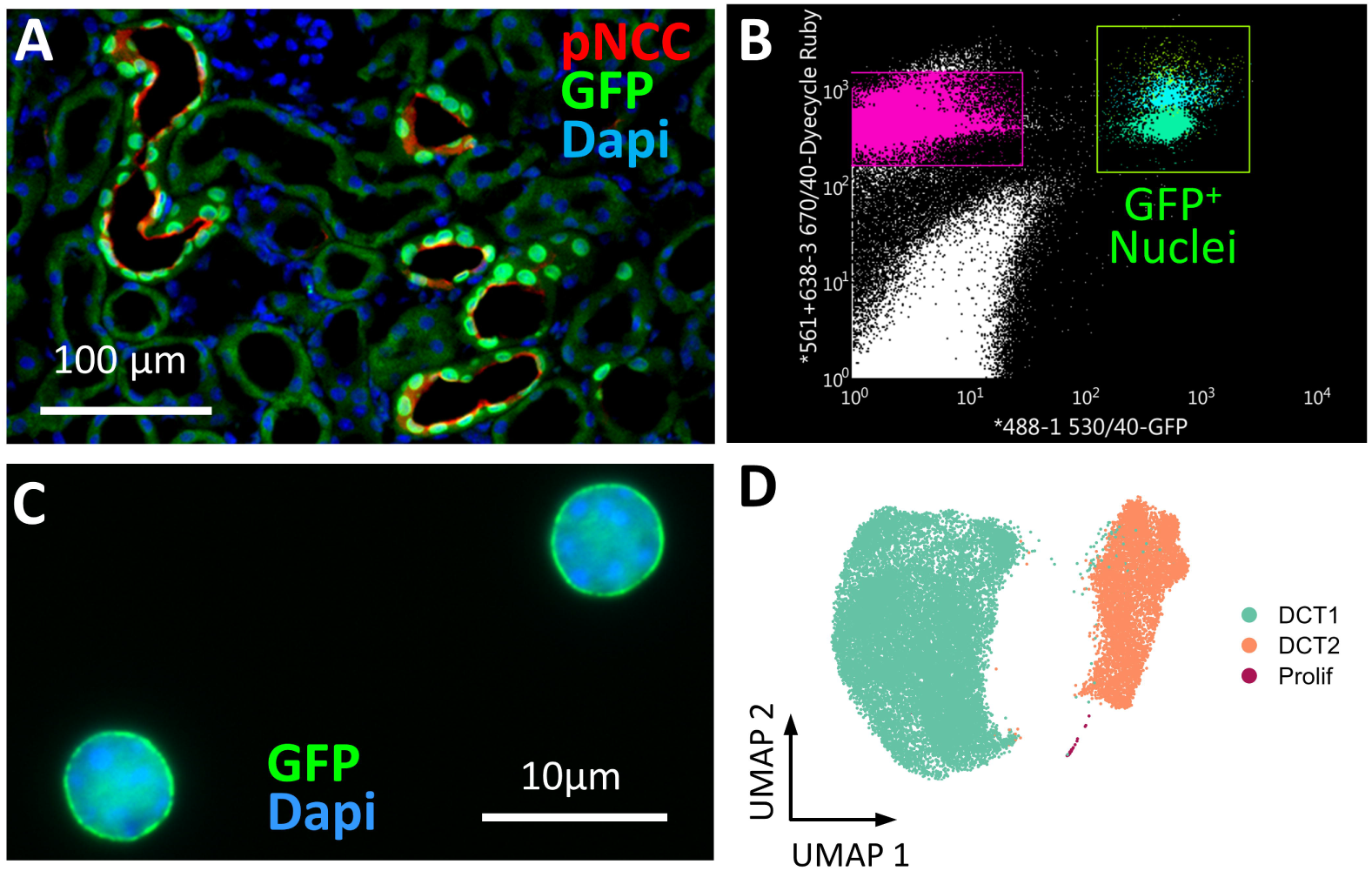
Enriched DCT snRNA-Seq confirms the DCT1 and DCT2 segmentation. **A)** Immunofluorescence image of the colocalization of pNCC (red) and INTACT (green). Nuclei is stained using DAPI in blue. **B)** Example of GFP^+^-nuclei sort gating. **C)** Image of sorted GFP^+^-nuclei at high magnification showing healthy nuclei with green INTACT (GFP) signal on the surface and blue DNA (DAPI) signal completely enclosed within the nuclei. **D)** UMAP projection for nuclei from three 10X Chromium control mouse DCT datasets. Different clusters are colored and annotated on the basis of **Figure S4**.

Targeting sequencing 10,000 nuclei per animal, led to average recovery of 12,513 nuclei/animal with an average of the median gene number per nucleus of 1843 and an average of mean reads per nucleus of 23,878 (**Supplemental table 2**). Following analysis through the Seurat pipeline that included doublet removal with DoubletFinder, ambient RNA removal with SoupX, normalization with scTransform, and integration with Seurat, we generated a 2D reduction of the first 20 principal components using uniform manifold approximation and projection (UMAP) to represent cellular clustering. The vast majority of nuclei expressed Slc12a3 (91.3%), encoding the canonical DCT marker NCC, the basis for the Cre-recombinase expression; however, there were additional populations that expressed markers that define other segments of the nephron in addition to Slc12a3 (**Figure S2**). Based on this incongruent marker expression and level of gene per nucleus, (**Figure S3**) we suspected that these were heterotypic doublet nuclei and filtered them before proceeding with the subsequent analysis.

After filtering suspected doublet populations, we observed 3 distinct populations of distal convoluted cells from DCT-INTACT mice that include DCT1, DCT2, and proliferating cells (**Figure 1D, S4**) similar to the distal tubule scRNA-Seq dataset that has previously been reported by Chen *et al.* ^9^. We directly compared the list of differentially expressed genes (DEGs) that define DCT1, DCT2 and proliferating cells between our nuclei-based strategy from their cell-based strategy and found a strong correlation between the same populations in both the enriched DCT snRNA-Seq dataset and the targeted distal tubule scRNA-Seq dataset (**Figure S5**) leading us to believe that these are the same populations of cells. In contrast to Chen *et al.*, we found the proportion of DCT1 to DCT2 to be 3:1 rather than 14:1 as they have reported. The ratio of 3:1 is similar to that measured by Welling and colleagues ^11^ using immunohistochemical techniques, suggesting that DCT-INTACT approach captures an accurate representation of the proportion of DCT1 and DCT2 within kidney tissue.

## 2) Functional and developmental heterogeneity along the DCT is revealed by enriched snRNA-Seq

The embryological origins of cells along the distal nephron have remained controversial, but single cell data from McMahon and colleagues suggested the existence of transitional cells ^12^. To explore the molecular heterogeneity of DCT1 and DCT2 populations and to provide information about its functional implications, we determined the top 10 DEGs by fold difference that distinguish DCT1 from DCT2 (**Figure 2A**). In contrast, the proliferating cell population has lower expression of these top 10 DEGs, but higher expression of cell-cycle machinery and proliferative genes, including *Top2a, Cenpp, Cep128, Knl1, Mki67,* etc. (**Figure 2A, Supplemental table 3**). We amalgamated the expression of the top 10 DEGs for DCT1 (*Erbb4, Egf, Trpm7, Fgf13, Col5a2, Umod, Ptgfr, Stk32b, Rtl4, Abca13*) and DCT2 (*Slc8a1, Arl15, Calb1, Slc2a9, Phactr1, Gls, S100g, Kl, Klk1, Egfem1)* using the “AddModuleScore” function in Seurat, to generate a DCT1 or DCT2 score. The visualization of this score shows clear distinction between DCT1 and DCT2 cells (**Figure 2B-C**). We also calculated the Pseudotime using Monocle 3; Pseudotime is a measure of how much progress an individual cell has made through a process such as cell differentiation ^42–45^. In many biological processes, cells do not progress in perfect synchrony, but transition from one state to the next according to their progress along a learned trajectory ^42–45^. Put simplistically, Pseudotime is the distance between a cell and the start of the trajectory. The trajectory within DCT is continuous starting within DCT1 and progressing to DCT2 (**Figure 2D**), suggesting that DCT2 is more differentiated than DCT1 or proliferating DCT cells. The expression of the DCT1 or DCT2 score-driving genes are plotted against Pseudotime as z-score (**Figure 2E-F**). These results indicate that even though DCT1 and DCT2 have separate functions, the transition between them is gradual.

**Figure 2.**
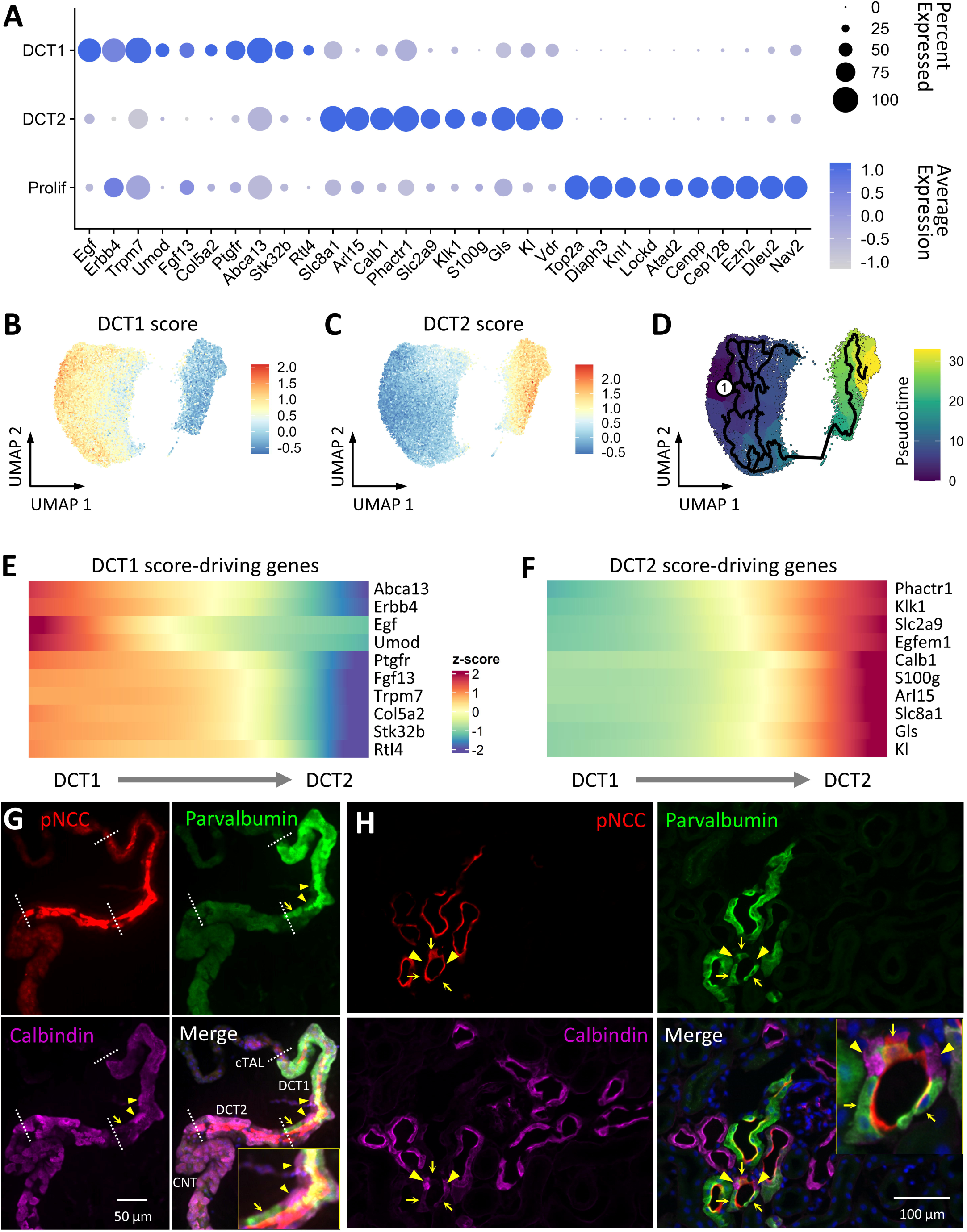
The transition between DCT1 and DCT2 is gradual. **A)** Dot plot showing the expression of top 10 DEGs in DCT1, DCT2 and proliferating cells across all clusters. Data are normalized and scaled (z-score) to examine relative expression across the cell clusters; “Average Expression” is the z-score of the average gene expression of all cells within a cluster (scaled values); and “Percent Expressed” is the percentage of cells with non-zero gene expression. **B-C)** UMAP projection for DCT1 **F)** and DCT2 **G)** score expression in all nuclei. The color indicates the expression of scores. **D)** UMAP projection of the Pseudotime. The “1” with a circle indicated the cell at time “0” or the “roots” of the trajectory. The black lines show the structure of the graph. DCT1 cells undergo less transcriptional change from the starting state to the end state; whereas DCT2 cells undergo more. **E-F)** A bar plot showing the z-scores of the DCT1 **I)** and DCT2 **J)** top 10 DEGs expression along the Pseudotime (DCT1 to DCT2). **G-H)** Immunofluorescence of micro-dissected tubule **K)** and thin sections **L)**. pNCC (red) labels the entire DCT; Parvalbumin (green) labels DCT1; Calbindin (magenta) labels DCT2 and CNT. Arrow heads point to DCT2 cells in the DCT1 and DCT2 transition segment; whereas arrows point to DCT1 cells in that segment.

While transcriptomics provides high resolution information about cell properties, it does not provide information about spatial cell location in the tubule segments. To study the transition between DCT1 and DCT2, immunostaining in micro-dissected tubules and thin sections was applied. The more proximal Thick Ascending Limb (TAL) and DCT have distinct morphological and molecular features; immunostaining in micro-dissected tubules showed a sharp border between TAL and DCT (**Figure 2G**). In contrast, immunofluorescence in micro-dissected tubules (**Figure 2G**) and thin sections (**Figure 2H**) revealed that DCT1 and DCT2 do not have a clear border. Parvalbumin (*Pvalb*, DCT1 marker) expression is high in DCT1 and gets weaker and spotty towards DCT2. In the “transition zone” between DCT1 and DCT2, parvalbumin and calbindin (DCT2 and CNT marker) expression is intermingled, confirming that the transition between DCT1 and DCT2 is gradual.

DCT2 (with the CNT) is the major site for active transcellular calcium reabsorption and exhibits electrogenic Na and K transport ^48, 49^. In contrast, magnesium transport has generally been described as occurring throughout the DCT ^50^. Our high-resolution transcriptomic data, however, indicate a separation between primary sites of calcium and magnesium handling, with more abundant magnesium handling genes along DCT1 (**Figure 3A, S6A**) and more abundant for calcium handling genes along DCT2 (**Figure 3B, S6B**). As previously suggested, NCC (*Slc12a3*)a nd its regulatory factors are more highly expressed by DCT1 cells, whereas both NCC and ENaC, as well as ENaC regulatory proteins are expressed along DCT2 (**Figure 3C, S6C**). Consistent with others’ reports ^9, 51, 52^, the alpha subunit of ENaC is widely expressed along the distal nephron, including DCT1 cells.

**Figure 3.**
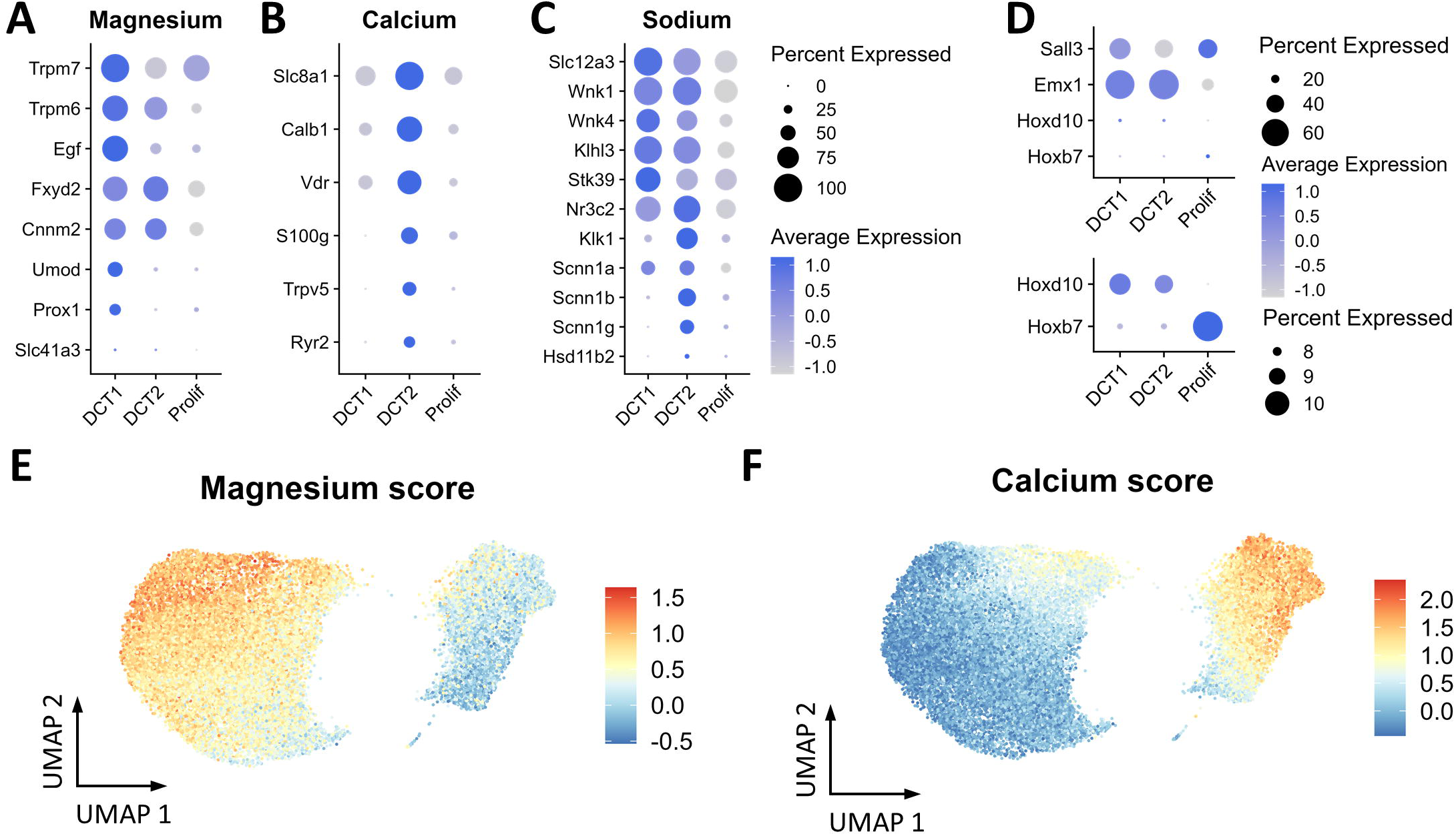
DCT1 and DCT2 have different expression patterns in electrolyte transport proteins. **A-C)** Distributions of transcripts associated with magnesium **A)**, calcium **B)** and sodium **C)** transport. Data are normalized and scaled (z-score) to examine relative expression across the cell clusters; “Average Expression” is the z-score of the average gene expression of all cells within a cluster (scaled values); and “Percent Expressed” is the percentage of cells with non-zero gene expression. **D)** Dot plot of *Hoxd10* (nephron progenitor origin homeobox gene) and *Hoxd7* (ureteric bud origin homeobox gene). **E-F)** UMAP projection for magnesium **E)** and calcium F) score expression in all nuclei. The color indicates the expression of the scores. **G-H)** DCT1 **G)** and DCT2 **H)** DEG pathway analysis. Significant cell-specific DEG from DCT1 or DCT2 were used to perform pathway analysis. The number of genes involved in each pathway is shown and the top 36 (DCT1) or 30 (DCT2) are highlighted. Some pathways are listed here and the full list is in **S9-10).**

To better characterize the expression patterns for genes involved in divalent ion metabolism, we defined a *magnesium cassette*, with genes relevant to magnesium transport (*Trpm6, Trpm7, Egf, Umod, Prox1, Fxyd2, Cnnm2* and *Slc41a3*) ^25–33^, and a *calcium cassette*, with genes relevant to calcium transport (*Slc8a1, Calb1, Vdr, S100g, Trpv5* and *Ryr2*) ^34–40^. As we did with the DCT1 and DCT2 scores, we amalgamated the magnesium/calcium cassette gene expression into magnesium and calcium scores. As expected, the magnesium cassette is enriched in DCT1 cells, whereas the calcium cassette is enriched in DCT2 cells (**Figure 3E-F**). The DCT1/DCT2/magnesium/calcium scores were shown to be consistent with scores derived for the enriched distal nephron scRNA-Seq dataset ^9^ (**Figure S7**).

The human distal nephron has not been characterized in the detail we have provided here. To compare mouse and human DCT cell transcriptomic profiles, we acquired the human snRNA-Seq data from the Kidney Precision Medicine Project (KPMP) ^53^. Data were downloaded on November 29^th^, 2022. This dataset includes 13 healthy reference, 10 chronic kidney disease (CKD) patients, and 6 acute kidney injury (AKI) patients. We separated DCT cells (DCT1, DCT2, cycling DCT and degenerative DCT as in the original KPMP nomenclature) from all cells (**Figure S8, 4A**) and applied the DCT1/DCT2/magnesium/calcium scores to the human DCT cells. Like in mice, DCT1 and magnesium scores are higher in human DCT1 cells than in DCT2 cells, and DCT2 and calcium scores are higher in DCT2 cells than in DCT1 cells (**Figure 4B-E**). We then compared the list of DEGs that define DCT1, DCT2 and proliferating cells between mouse and human DCT nuclei and found a significant correlation between DCT1, DCT2 and the proliferating cell (cycling DCT in KPMP human DCT cells) markers (**Figure 4F-G**), which confirms that human and mouse DCT segmental functions are organized similarly.

**Figure 4.**
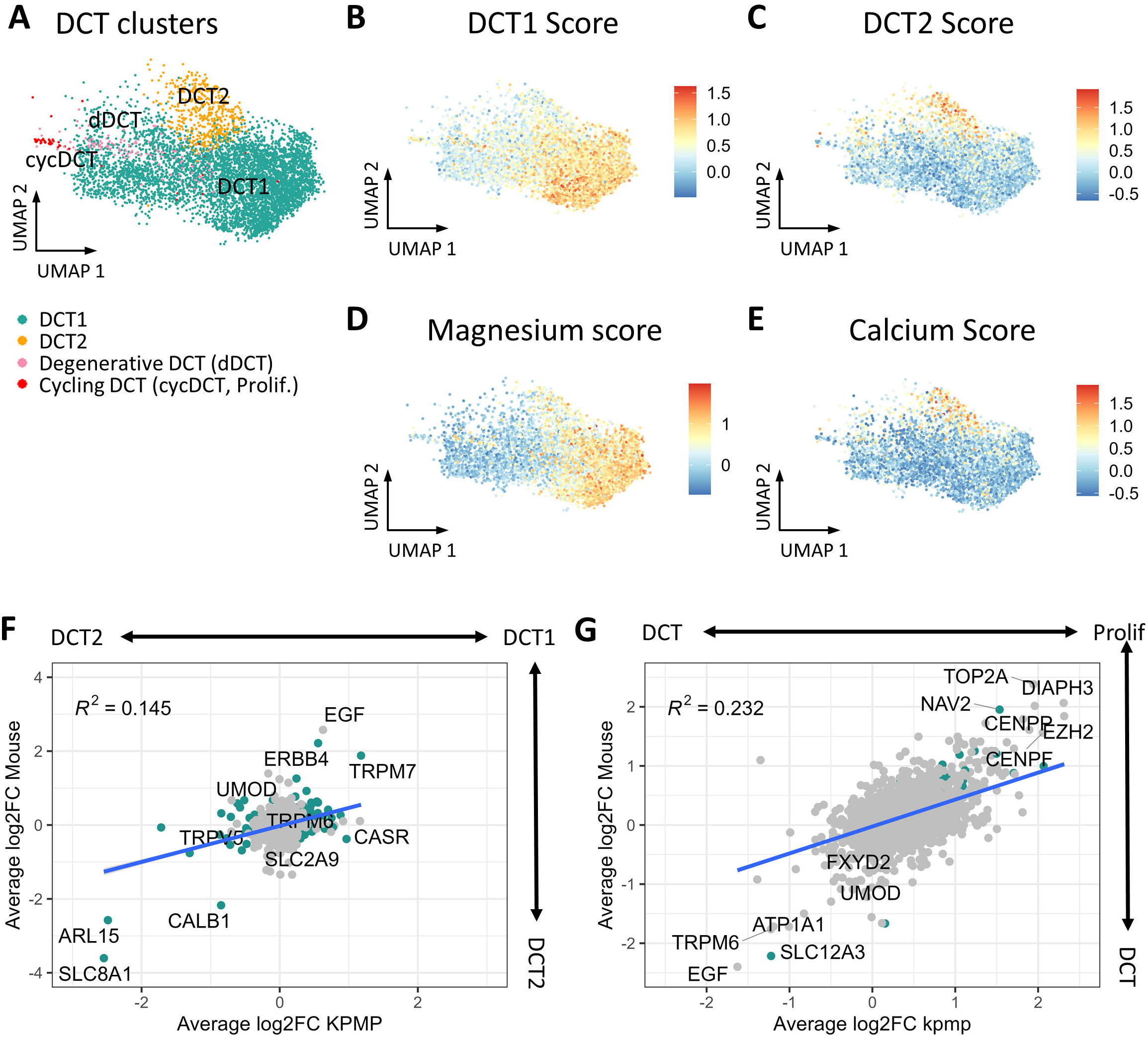
Human DCT cells from Kidney Precision Medicine Project (KPMP) dataset have similar signatures to mouse DCT cells. **A)** UMAP projection for all DCT cells from KPMP snRNA-Seq dataset. The annotations are originally from KPMP. The dDCT is degenerative DCT; cysDCT is cycling DCT, which is similar to the proliferating DCT cells in our dataset. **B-E)** UMAP projection for DCT1 **B)**, DCT2 **C)**, magnesium **D)** and calcium **E)** score expression in all DCT nuclei. The color indicates the expression of the scores. **F-G)** Correlation of cluster-defining DEG between mouse and human DCT1 versus DCT2 **F)** and DCT versus proliferating cells **(G)**. The x-axis is the average log2 fold change of the human DEG; y-axis is the average log2 fold change of the mouse DEG. R^2^ is the coefficient of determination. The DEGs with adjusted p value less than 0.01 in both mouse and human are highlighted.

The origin of cell types along the distal nephron has been controversial. Recent lineage tracing approaches suggest that DCT2 cells may derive from the ureteric bud, even though they express canonical markers of more proximal cells ^12, 54^. To explore this, we applied pathway analysis, which is a computational method that determines whether a priori defined set of genes show statistically significant differences ^46^. The pathway analysis revealed enrichment for well-known DCT1-related pathways, including metal ion transport, Na transport, protein autophosphorylation, and nutrient level response (**Figure 5A, S9**). Interestingly, and in stark contrast with DCT1, transcript enrichment in DCT2 cells predominantly involves renal system development, urogenital system development, and branching morphogenesis of epithelial tubules. (**Figure 5B, S10**). The functional importance of these differences will be discussed below.

**Figure 5.**
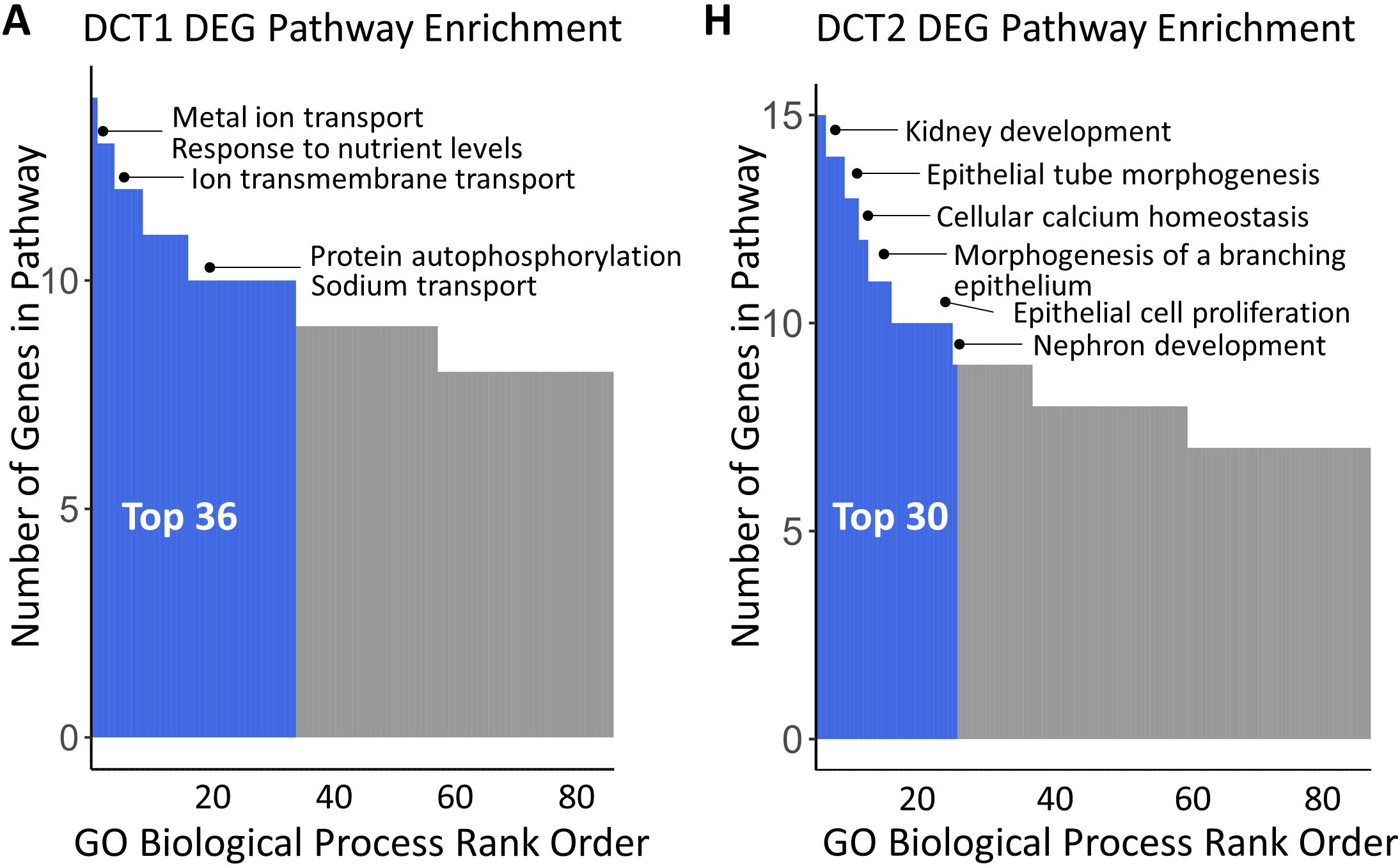
Transcripts enriched in DCT1 or DCT2 cells involve distinct pathways. **A-B)** DCT1 **A)** and DCT2 **B)** DEG pathway analysis. Significant cell-specific DEG from DCT1 or DCT2 were used to perform pathway analysis. The number of genes involved in each pathway is shown and the top 36 (DCT1) or 30 (DCT2) are highlighted. Some pathways are listed here and the full list is in **S9-10).**

## 3) Cell type transitions along the distal convoluted tubule are gradual

Given the postulated dichotomous origins of the DCT and CNT, cell types in these segments have been inferred to be discrete and, at segment junctions, mosaic ^12^. Yet, transcriptomic work in the brain and other tissues has shown that cell types may transition gradually from one to the other, complicating classification ^55^. To unmask additional layers of cell type heterogeneity within each population, we separated DCT1 and DCT2 from the mouse DCT dataset (**Figure 1D**) and re-clustered at higher resolution. This analysis confirmed as many as four populations (DCT1 A-D) ^9^ of DCT1 cells can be identified (**Figure 6A, S11**). Yet, our high-fidelity transcriptomics showed that, instead of being distinct cell types, these populations express the same core set of genes and are discriminated based on gene expression gradients (**Figure 6B**). The cells, therefore, transition from more strictly DCT1-like in character to more strictly DCT2-like in character, with DCT1-A having the greatest abundance of DCT1 markers (*Egf* and *Erbb4)*, and DCT1-B having the greatest abundance of DCT2 markers (*Slc8a1* and *Calb1)*. For magnesium and Na handling genes, DCT1-A to −D has similar expression patterns, with higher expression of magnesium genes and DCT1-B has greater expression of calcium handling proteins; the latter is consistent with the top genes being similar to DCT2 (**Figure 6C, S13)**.

**Figure 6.**
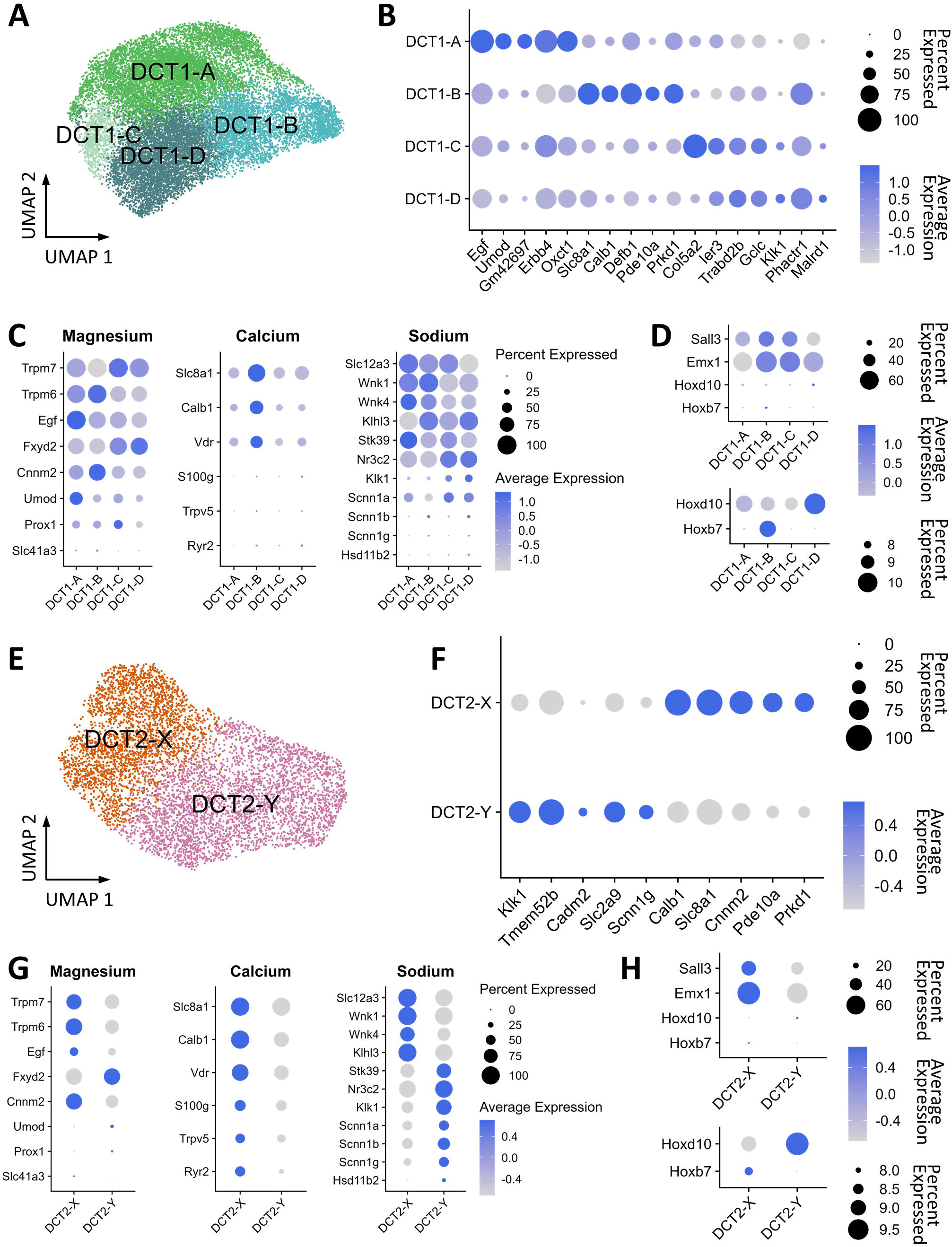
DCT1 cells are one cell type and DCT2 cells have two subtypes. **A)** UMPA projection for DCT1 subtypes within all DCT1 nuclei. **B)** Dot plot of top 5 genes expressed in DCT1A-D. **B)** Distributions of transcripts associated with magnesium, calcium and sodium transport. **C)** Dot plot of *Hoxd10* (nephron progenitor origin homeobox gene) and *Hoxd7* (ureteric bud origin homeobox gene) in DCT1A-D. **D)** UMPA projection for DCT2 subtypes within all DCT2 nuclei. **F)** Dot plot of top 5 genes expressed in DCT2-X and -Y. **G)** Distributions of transcripts associated with magnesium, calcium and sodium transport. **H)** Dot plot of *Hoxd10* (nephron progenitor origin homeobox gene) and *Hoxd7* (ureteric bud origin homeobox gene) in DCT2-X and -Y. Data are normalized and scaled (z-score) to examine relative expression across the cell clusters; “Average Expression” is the z-score of the average gene expression of all cells within a cluster (scaled values); and “Percent Expressed” is the percentage of cells with non-zero gene expression.

In contrast to the cell type gradients in DCT1, DCT2 cells comprised two patterns, DCT2-X and DCT2-Y. DCT2-X are enriched in NCC, magnesium, and calcium-related genes (including *Slc8a1* and *Calb1)*, and DCT2-Y is enriched in genes supporting electrogenic Na transport (including *Klk1* and *Scnn1g*) (**Figure 6E-F, S12**). These patterns indicate that DCT2-X is involved in divalent cation transport, whereas DCT2-Y, which is enriched in ENaC (*Scnn1a*), 11-β-dehydrogenase isozyme 2 (*Hsd11b2*), mineralocorticoid receptor (*Nr3c2*), and Kallikrein-1 (*Klk1*), mediates electrogenic Na transport (**Figure 6G, S13**). As snRNA-Seq does not provide spatial data, and our immunohistochemistry cannot distinguish cell subtypes, we cannot conclude whether the DCT2-X and DCT2-Y cells are intermingled or anatomically sequential.

Homeobox genes have been considered to be major drivers of morphological development in animals ^56, 57^. The expression patterns of the 39 mammalian homeobox genes are distinct in DCT1, DCT2, and proliferating cells (**Figure S14**), suggesting that they develop along different trajectories. Within the homeobox gene family, *Hoxb7* is specific to the ureteric bud derivatives, and *Hoxd10* is specific to the cells of nephron progenitor origin ^12^. The expression of *Hoxb7* in proliferating cells is the highest, and *Hoxd10* highest in DCT1 (**Figure 3D**). *Emx1* has been reported to be an important regulator of distal tubule development in zebrafish ^58^, which is not selective for DCT1 comparing to DCT2; whereas *Sall3* is a DCT-selective zinc-finger protein ^59^, which is higher in proliferating cells and DCT1 compared to DCT2 (**Figure 3D**). Among the subpopulations, the expression of *Hoxb7* in DCT1-B is the highest (only lower than the proliferating cells), and *Hoxd10* higher in DCT1-D (**Figure 6D, 6H, Figure S14C**). Collectively, it suggests that DCT1 and DCT2 cells have mixed origins, some originate from the ureteric bud and some from the nephron progenitor. Yet, the DCT proliferating cells originate from ureteric buds.

## Discussion

The DCT-Cre INTACT approach allowed us to examine the distal nephron in unprecedented detail on cell types and cell states. In rodents and humans, the transition from DCT1 to CNT is gradual at the microscopic level ^60^; this is in contrast to the abrupt change from thick ascending limb to DCT. We observed this gradual transition in immunofluorescent experiments, but also noted that DCT1 and DCT2 cells are sometimes intermingled. Yang and colleagues ^4^ used similar techniques to categorize transitions between segments and suggested that the overlap between NCC and ENaC was minimal. To obtain more information about the nature of these transitional segments, we utilized snRNA-Seq. Zeng ^55^ recently described a comprehensive functional-transcriptomic analysis of brain cells, providing a useful definition of cells types and cell states. One feature that was noted was the coexistence of both discrete and continuous variations between cell types. Because we achieved a highly enriched population of DCT cells using the INTACT approach, we were able to determine that DCT1 cells and DCT2 cells are discrete, but are connected by a continuum of cell states (**Figure 2E-F).** This revealed heterogeneity of segments that are difficult to study using electrophysiological approaches for technical reasons, and suggested the existence of multiple cell states along this segment. The transition from DCT to CNT could not be evaluated here, as we used the *slc12a3* promoter to drive expression of the INTACT protein.

The current results provide important insights into the origin of the DCT and CNT, a subject of continuing controversy. Nephrons develop from mesenchymal progenitors and connect to the ureteric epithelial collecting system, which is developed from the ureteric bud ^61^. While most studies conclude that the CNT derives from the ureteric bud and DCT cells from mesenchymal progenitors, the nature of the transitional segment, comprising the DCT2 and ‘early CNT’ in rodents and humans, has been obscure ^62^. McMahon and colleagues combined fate mapping and transcriptomics to clarify this issue. They reported that both principal-like cells and intercalated cells in the CNT had both nephron and ureteric features ^61^.

Our work shows that cells gradually transition from DCT1 to DCT2 and likely CNT at the transcript level, supporting McMahon’s observation that these segments do not derive from one or the other embryological origin. Yet, although both DCT1 and DCT2 express more *Hoxd10* compared with *Hoxb7,* suggesting that most of DCT1 and DCT2 cells develop from nephron progenitors. The Pseudotime analysis discussed above suggested that DCT2 cells are more differentiated than DCT1 cells, indicating that DCT2 cells arise at a later time point during development. DCT2 cells are rich in transcripts related to kidney development, whereas DCT1 cells are rich in transcripts related to transport. Interestingly, despite being intermingled among DCT1 and DCT2 cells (immunofluorescence data not shown), the proliferating cells have more *Hoxb7,* suggesting that they develop from ureteric bud. As the complexity of DCT cell origin, it is speculated that the DCT is where the ureteric bud and the nephron join.

In summary, the nucleus isolation approach eliminates differences in recovery efficiency across cell types, thereby providing a more accurate representation of a small population of kidney cell. With the enriched cell type analysis, we will have the ability to generate high-fidelity resolution data with better clarity. Moreover, our-high resolution DCT transcriptome dataset can be used as a reference atlas for future work to map and identify DCT1, DCT2 and proliferating cells from non-enriched kidney nuclei datasets.

## Acknowledgement

We thank the OHSU Flow Cytometry, Gene Profiling, and Massively Parallel Sequencing core facilities for technical support.

## Table of Contents for Supplementary Information

Supplemental table 1

Supplemental table 2

Supplemental table 3

Supplemental Figures

