## Supplemental Data for "Enriched Single-Nucleus RNA-Sequencing reveals unique attributes of distal convoluted tubule cells"

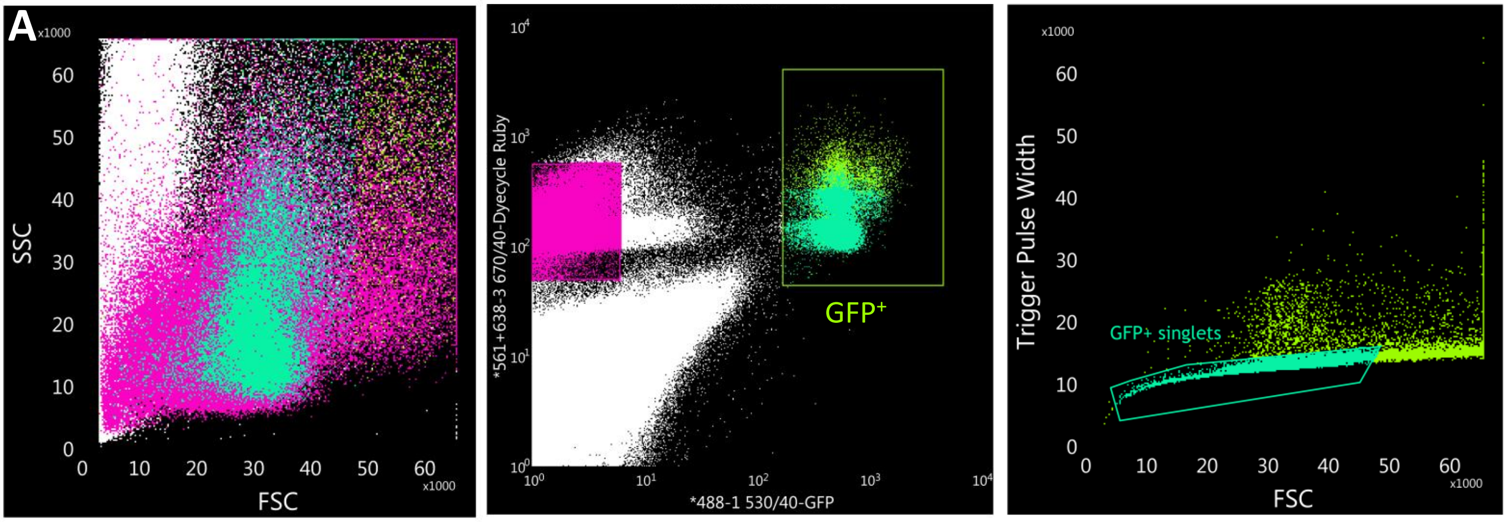


**B**

| Populations | Events | % Total | % Parent | FSC Mean | SSC Mean |
|---|---|---|---|---|---|
| All Events | 1,000,000 | 100.00% | #### | 12,622 | 22,575 |
| GFP+ | 24,617 | 2.46% | 2.46% | 37,264 | 29,068 |
| GFP+ singlets | 18,532 | 1.85% | 75.28% | 31,330 | 23,262 |
| GFP- | 250,780 | 25.08% | 25.08% | 28,253 | 19,801 |

**Figure S1. Gating strategy for DCT nuclei sorting.** A) Example plots show the nuclei were sorted against both GFP+(x-axis) and Ruby+(y-axis) in the middle panel; and against singlets in the right panel. B) A table summarizes the proportion of the nuclei after each gating from the example above.

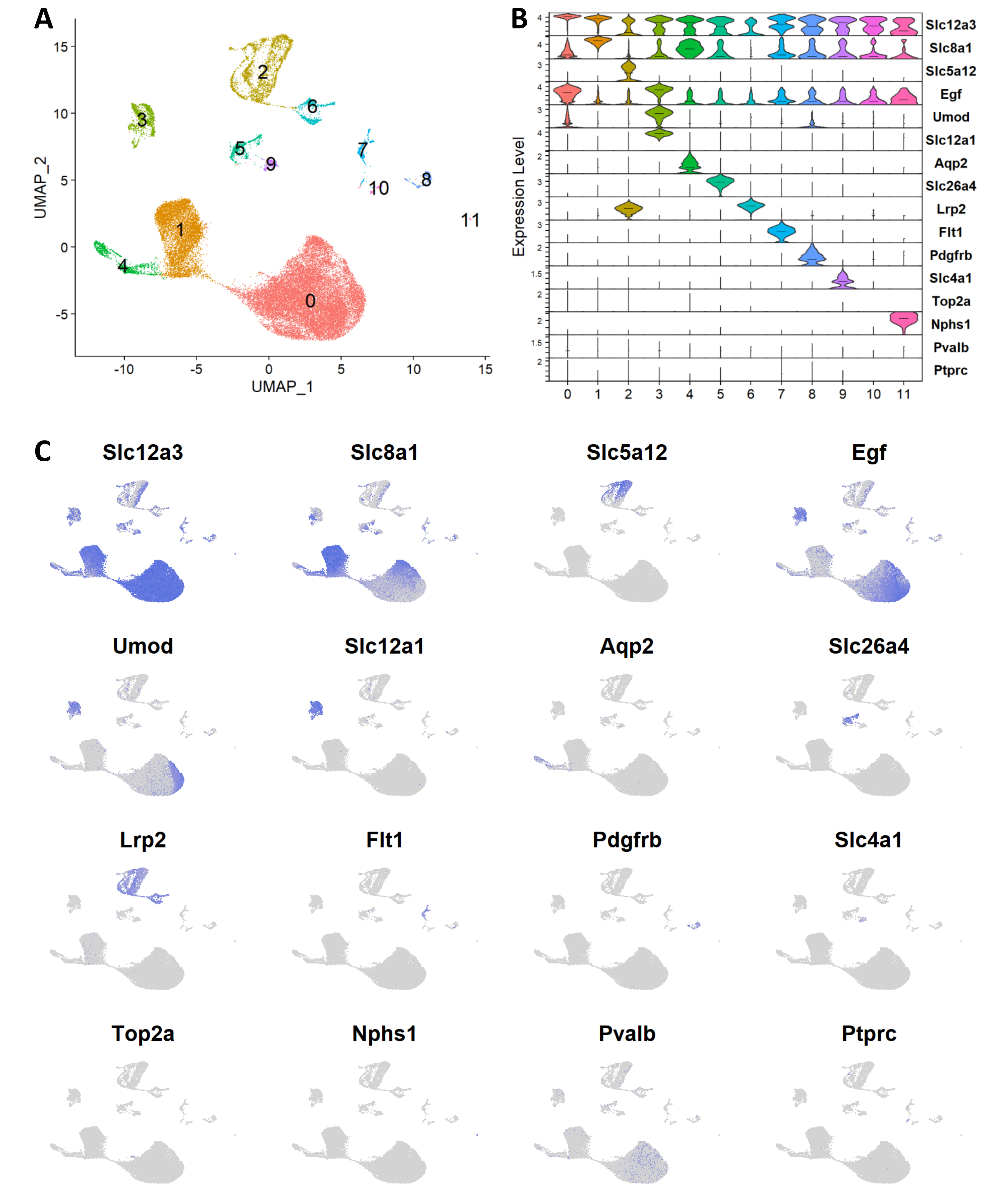


**Figure S2. Canonical markers expression for all cell type.** A) UMAP projection for all nuclei after initial pre-process and filtering. B) Violin plots for canonical markers of different cell type. C) UMAP projection for canonical markers.

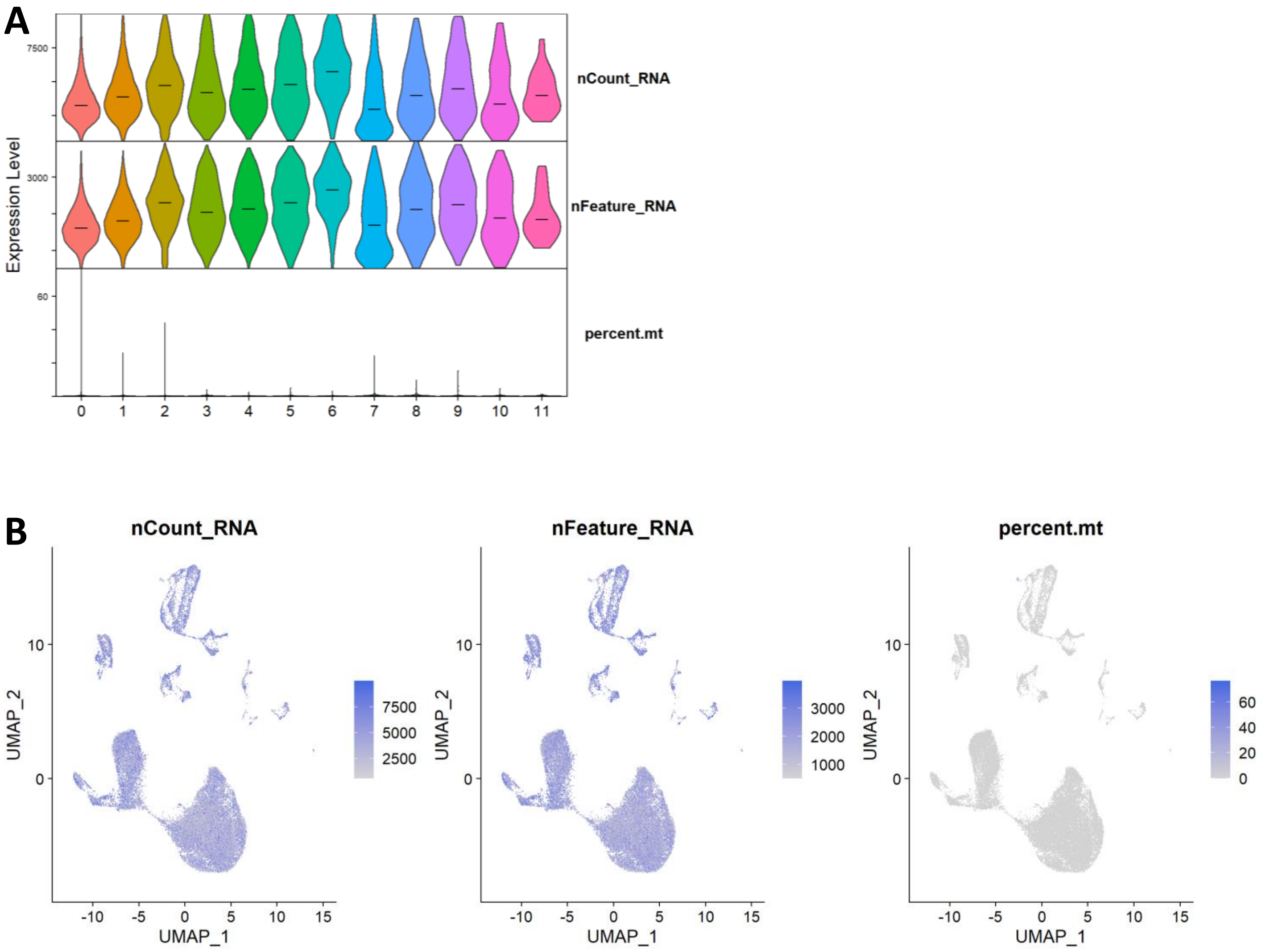


**Figure S3. Quality control of all nuclei.** A) Violin plots for nCount_RNA (number of RNA counts), nFeature_RNA (number of features), and percent.mt (mitochondria RNA percentage). B) UMAP projection for nCount_RNA, nFeature_RNA, and percent.mt

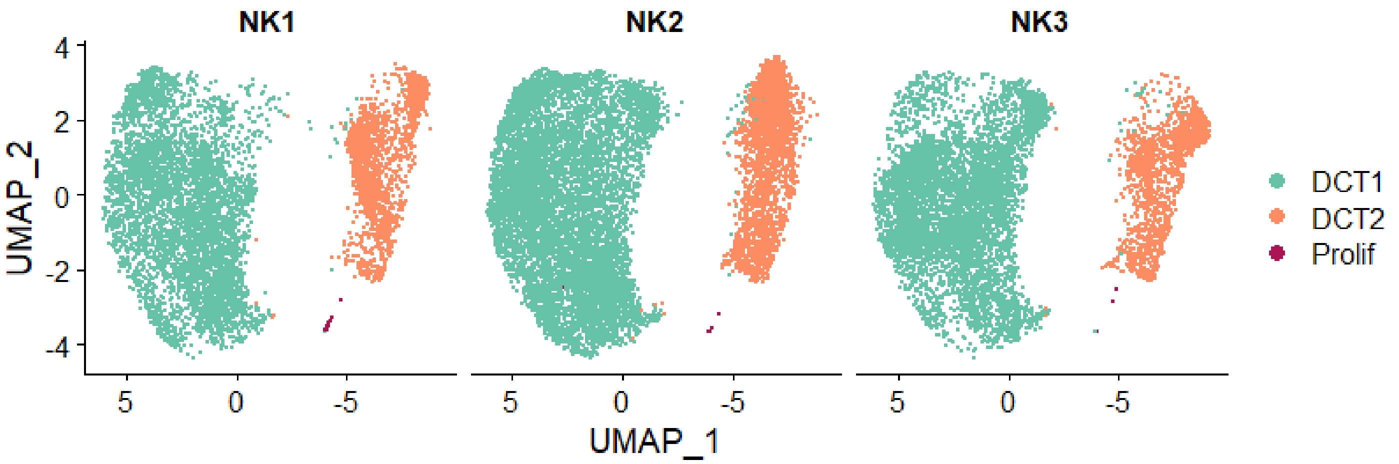


**Figure S4. Clustering of DCT nuclei in three control samples.** UMAP projection for nuclei from three 10X Chromium control mouse DCT datasets (NK1, NK2 and NK3). The distribution and clustering of all three samples are similar.

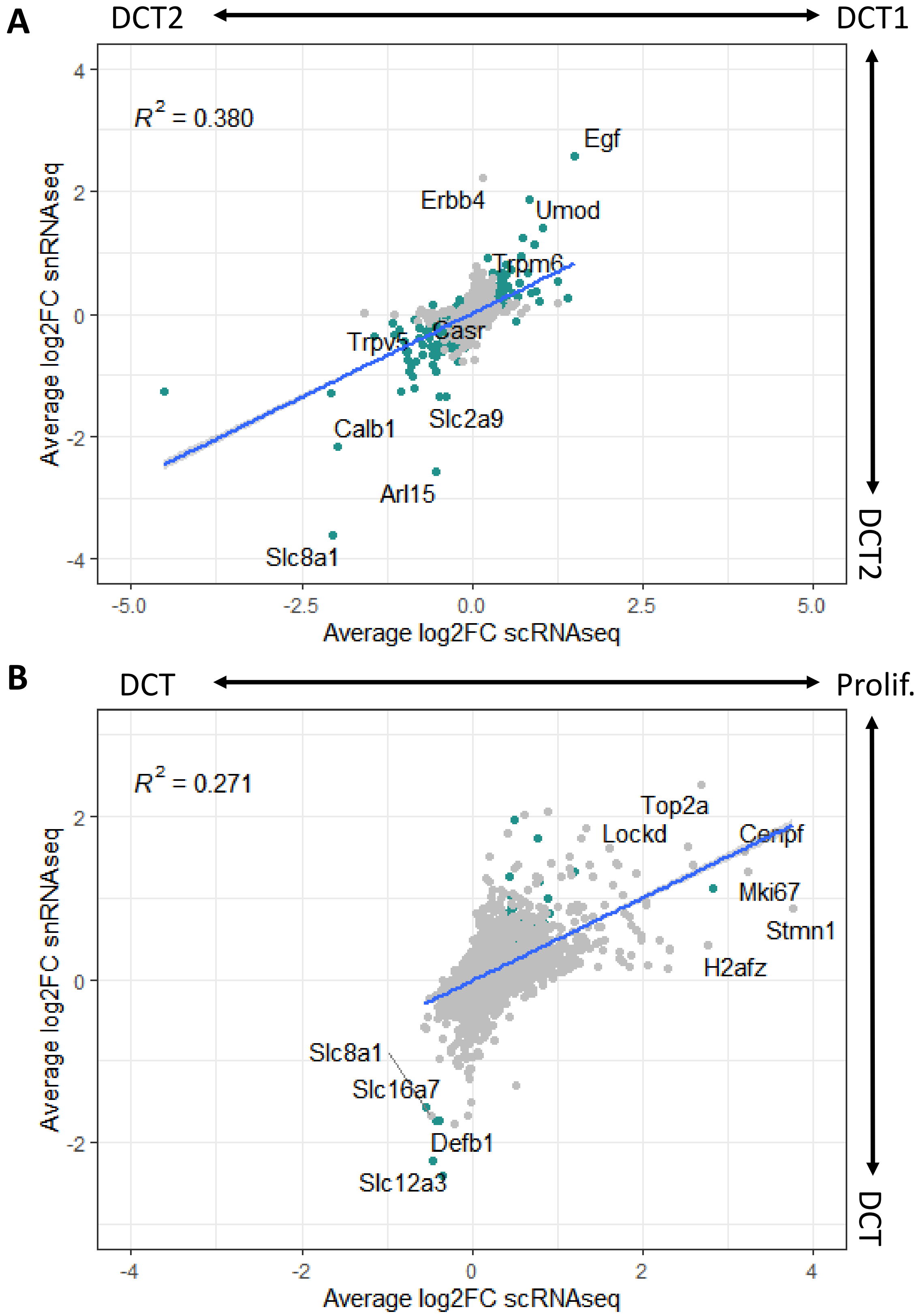


**Figure S5. Comparing snRNA-Seq data and previously published scRNAseq data of DCT cells.** A-B) Correlation of cluster-defining DEG between scRNAseq and snRNA-Seq DCT1 versus DCT2 A) and DCT versus proliferating cells B). The x-axis is the average log2 fold change of the scRNAseq DEG; y-axis is the average log2 fold change of the snRNA-Seq DEG. R2 is the coefficient of determination. The DEGs with adjusted p value less than 0.01 in both scRNAseq data and snRNA-Seq data are highlighted.

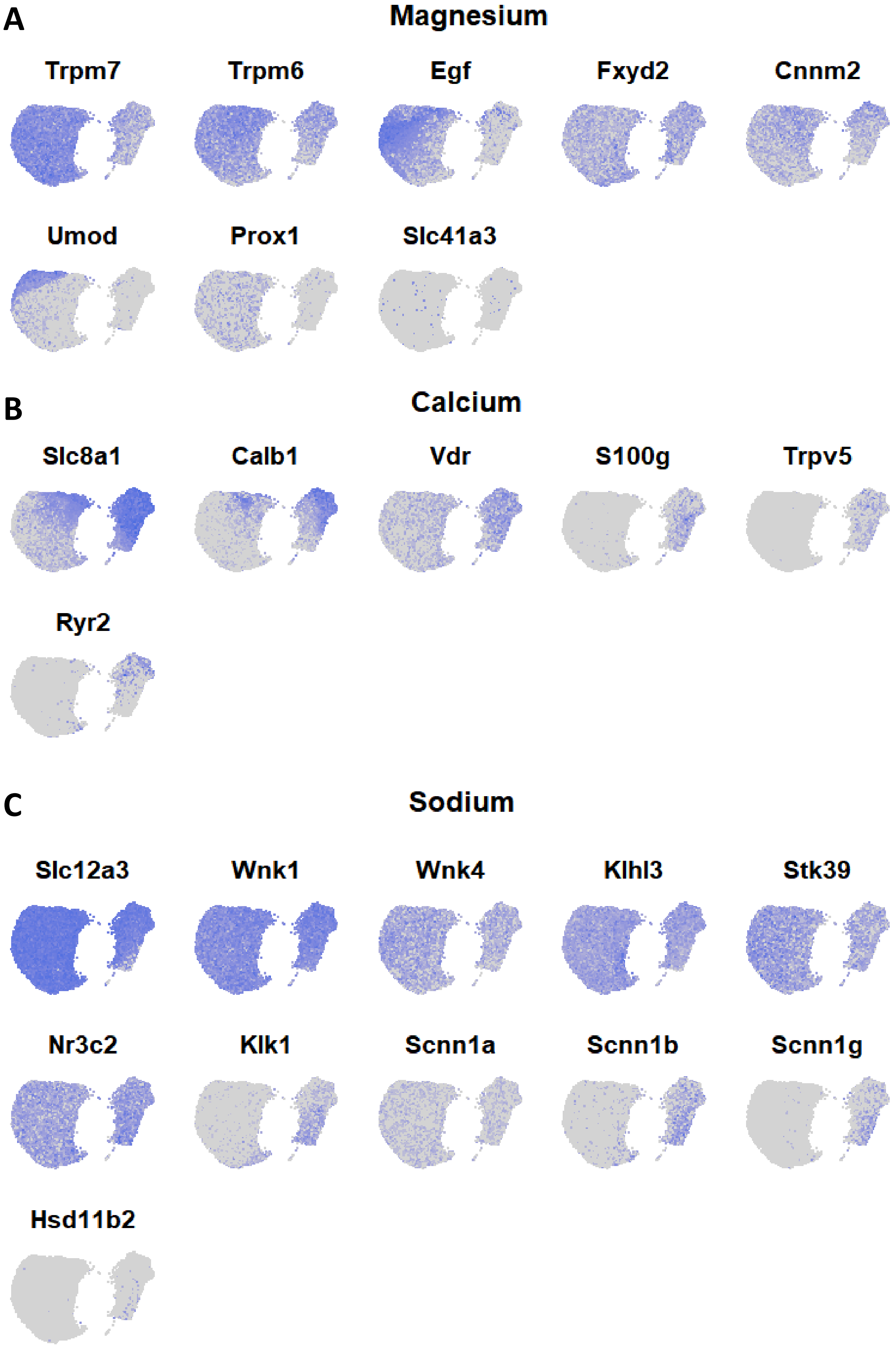


**Figure S6. Comparing snRNA-Seq data and previously published scRNAseq data of DCT cells.** A-C) UMAP projection for transcripts associated with magnesium A), calcium B) and sodium C) transport.

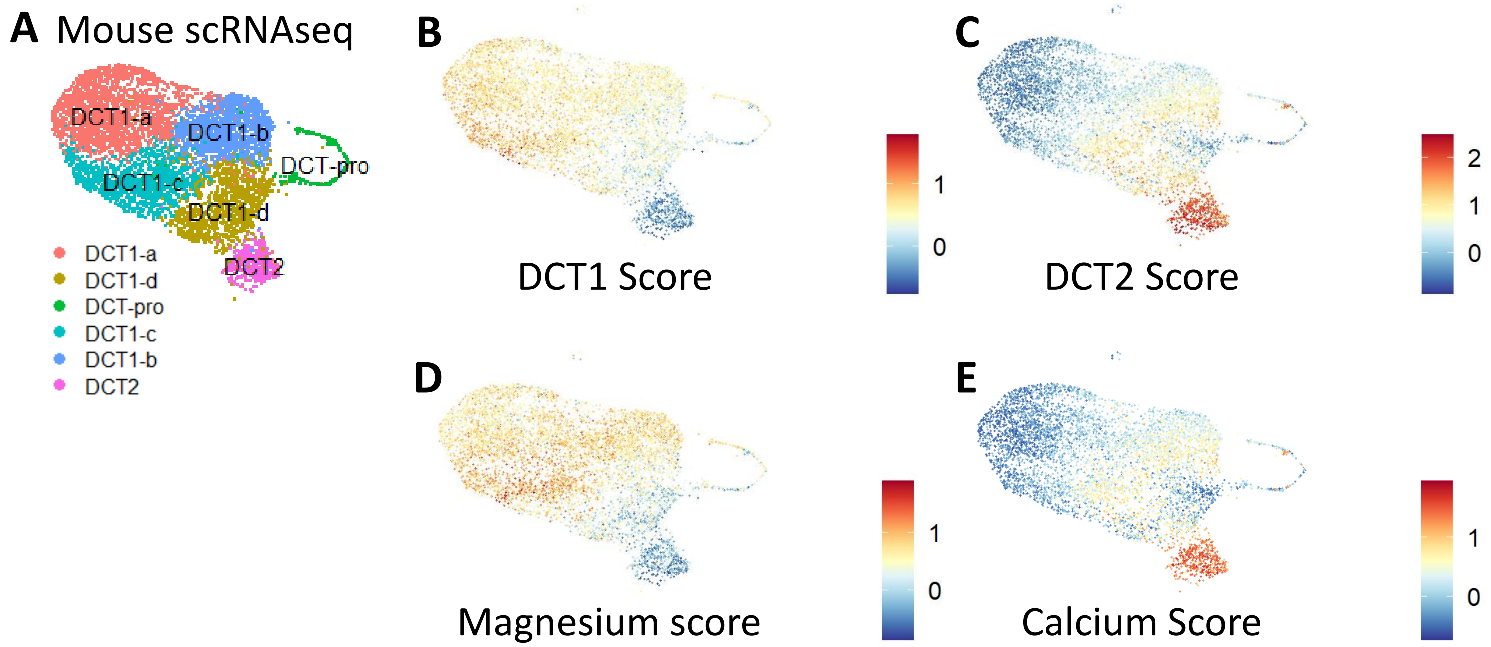


**Figure S7. Validating the scores in a previously published mouse scRNAseq dataset.** A) UMAP projection for DCT cells from a previously published mouse scRNAseq dataset. The annotations are from the original publication. DCT-pro is the proliferating cell population. DCT1 has DCT1a-d subtypes. B-E) UMAP projection for DCT1 B), DCT2 C), magnesium D) and calcium E) score expression in all DCT nuclei. The color indicates the expression of the scores.

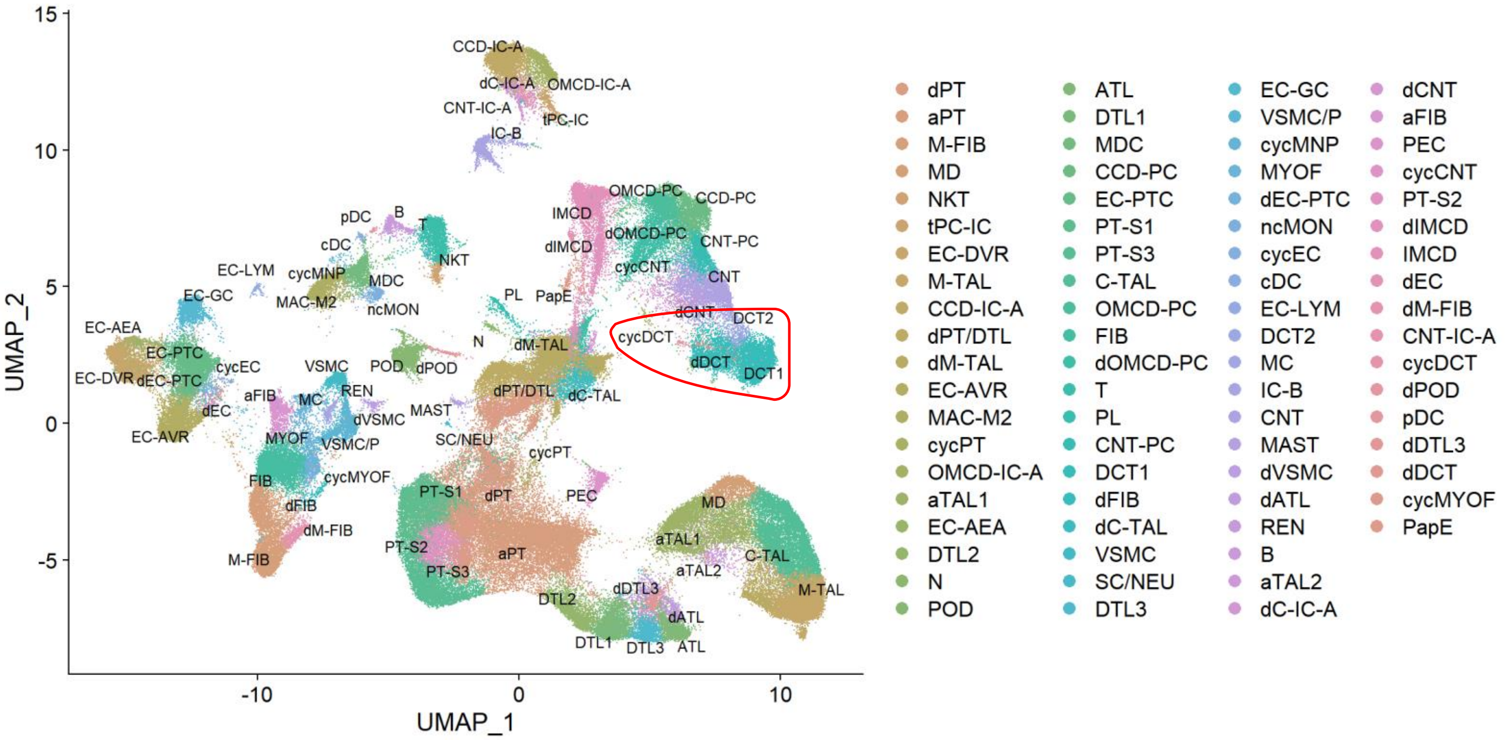


**Figure S8. The Kidney Precision Medicine Project (KPMP) snRNA-Seq dataset.** UMAP projection for all cells from the KPMP snRNA-Seq dataset. The annotations are from the original dataset. The DCT clusters (in red circle) were separated for the comparison between human and mouse DCT cells in Figure 5.

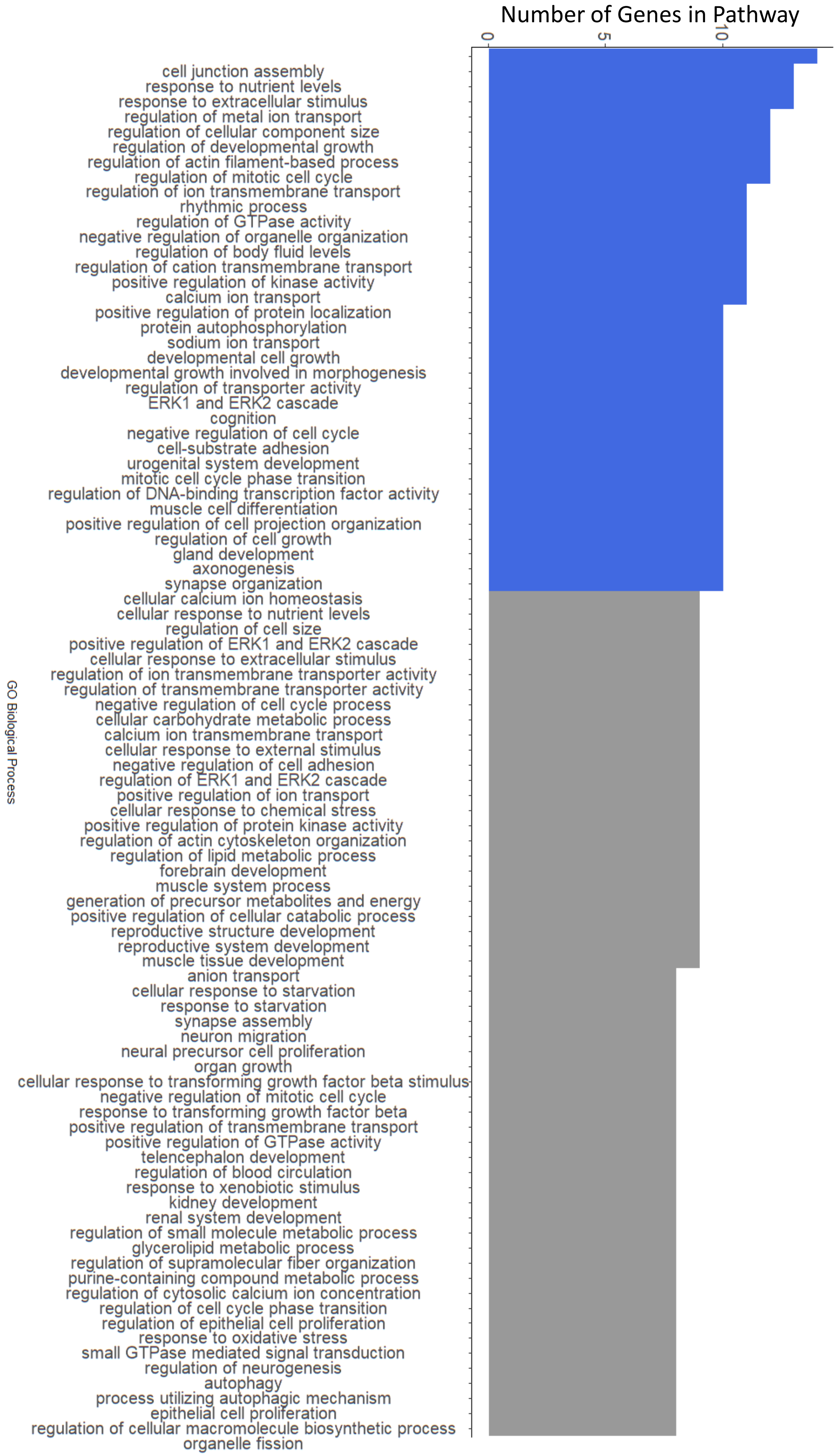


**Figure S9. DCT1 DEG pathway analysis.** Significant cell-specific DEGs from DCT1 were used to perform pathway analysis. The number of genes involved in each pathway is shown and the top 36 are highlighted.

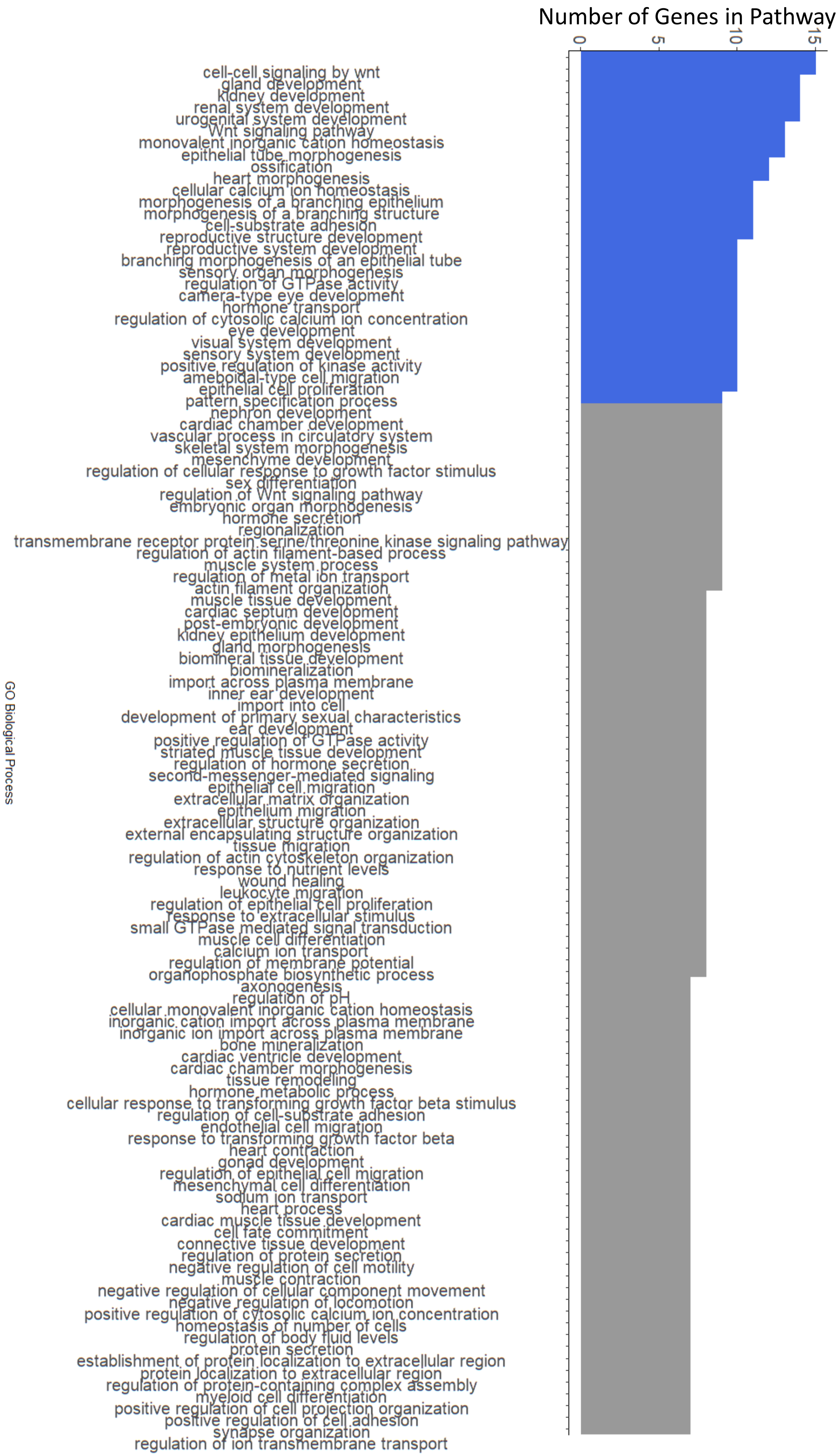


**Figure S10. DCT2 DEG pathway analysis.** Significant cell-specific DEGs from DCT2 were used to perform pathway analysis. The number of genes involved in each pathway is shown and the top 30 are highlighted.

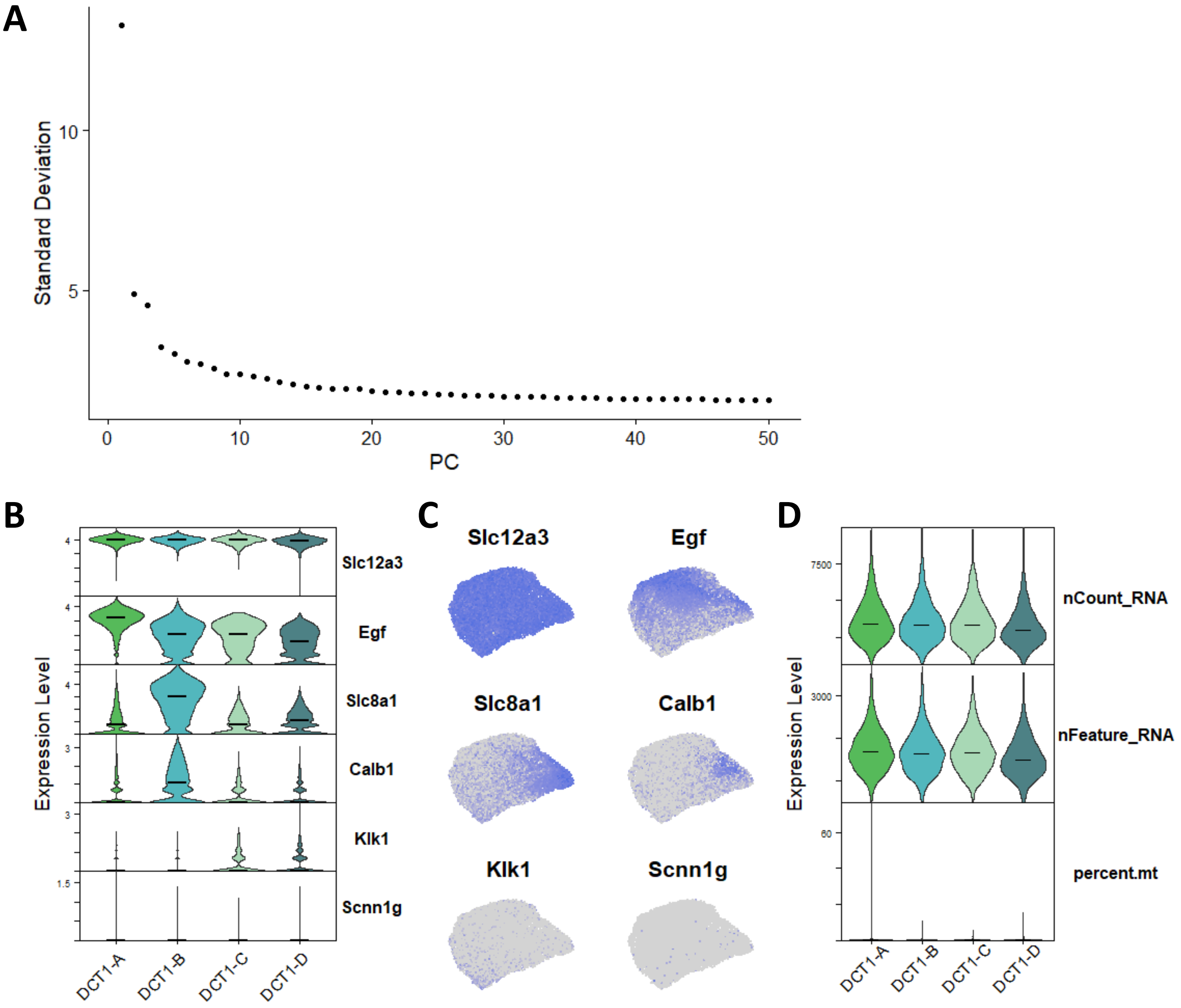


**Figure S11. DCT1 sub-population.** A) Principal component analysis of DCT1 cells. B-C) Violin plots B) UMAP projection C) of canonical markers in DCT1A-D. D) Violin plots for nCount_RNA (number of RNA counts), nFeature_RNA (number of features), and percent.mt (mitochondria RNA percentage) in DCT1A-D.

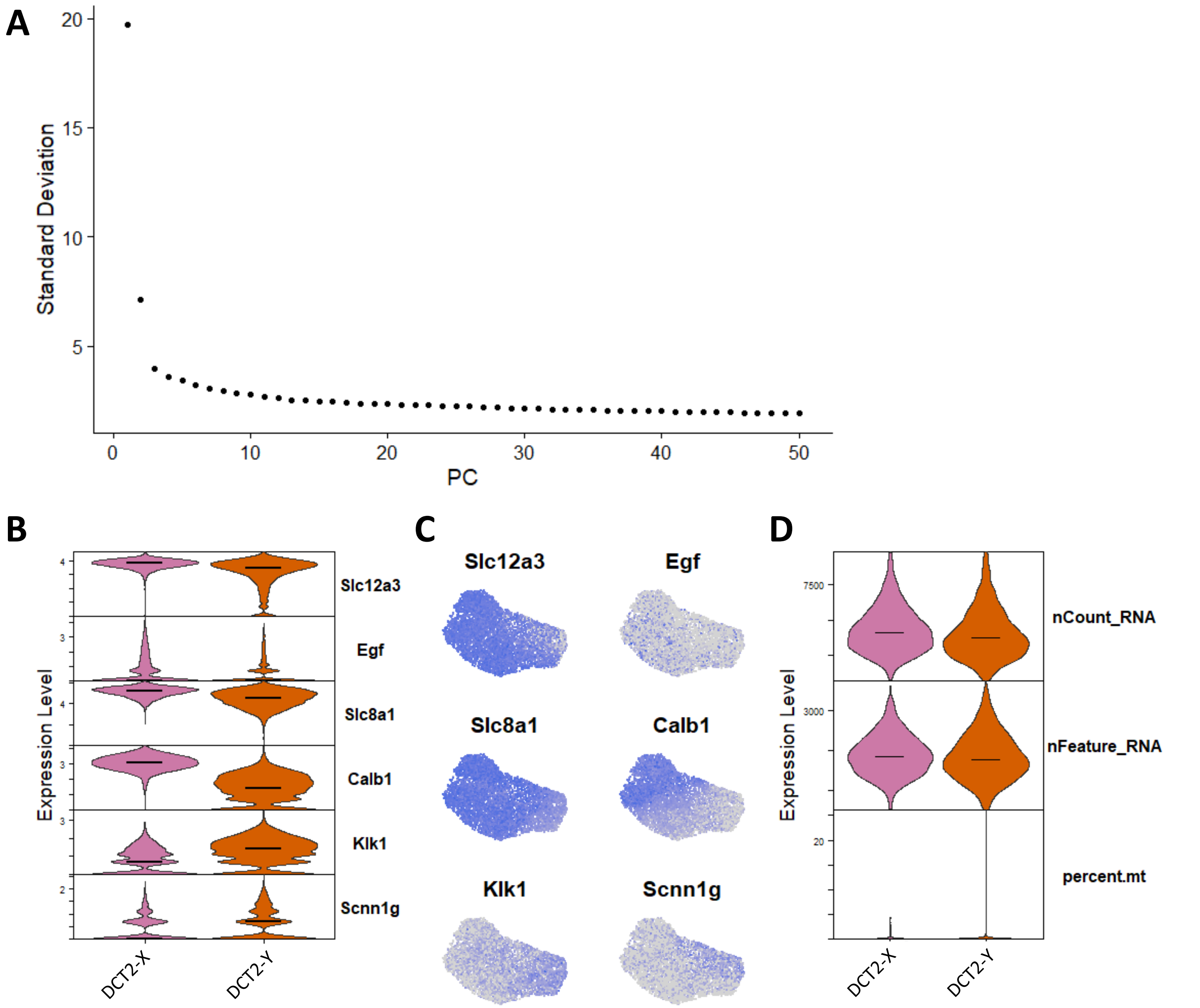


**Figure S12. DCT2 sub-population.** A) Principal component analysis of DCT2 cells. B-C) Violin plots B) UMAP projection C) of canonical markers in DCT2X-Y. D) Violin plots for nCount_RNA (number of RNA counts), nFeature_RNA (number of features), and percent.mt (mitochondria RNA percentage) in DCT2X-Y.

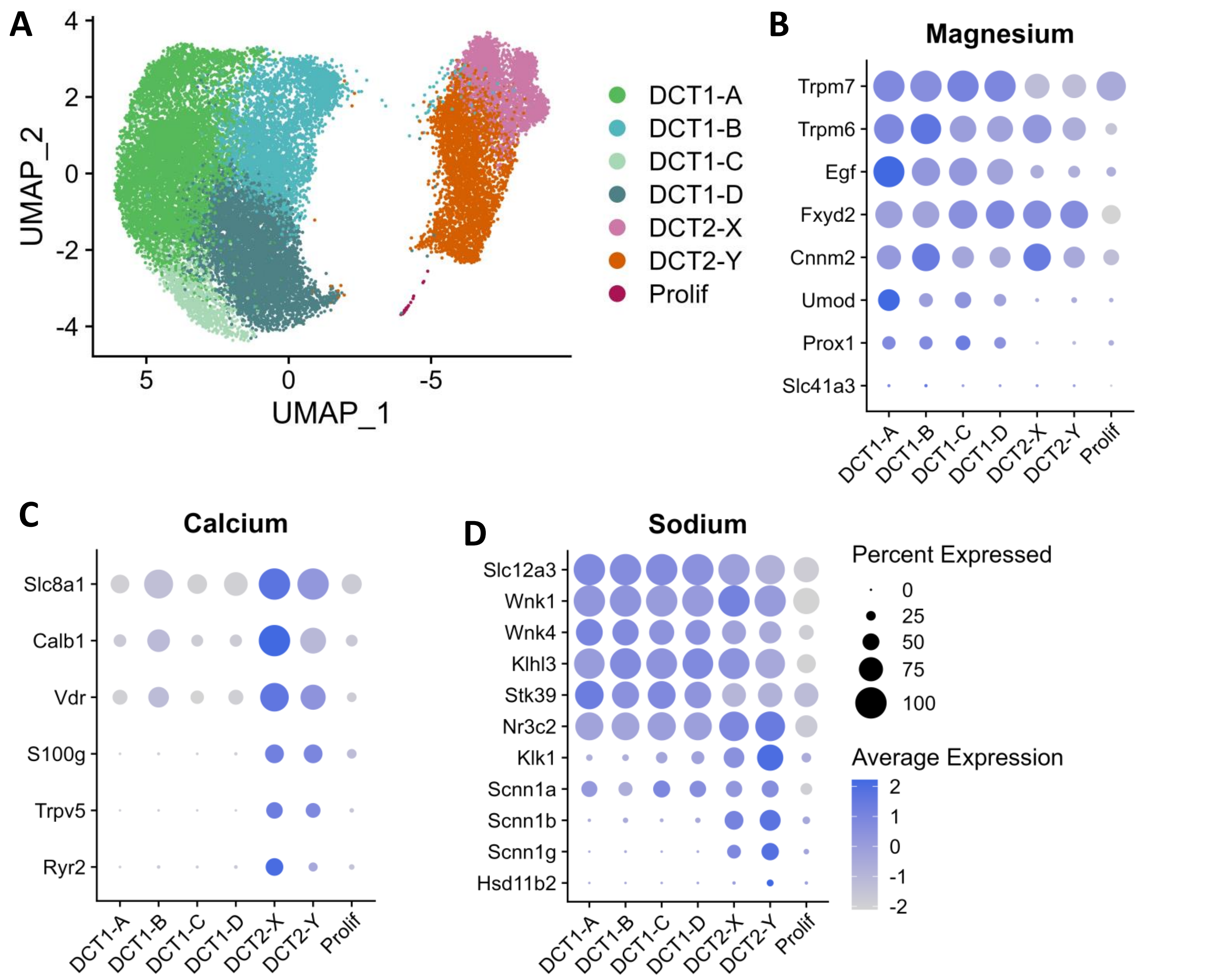


**Figure S13. Distributions of transcripts associated with magnesium, calcium and sodium transport.** A) UMPA projection for all DCT subtypes within all DCT nuclei. B-D) Distributions of transcripts associated with magnesium B), calcium C) and sodium D) transport. Data are normalized and scaled (z-score) to examine relative expression across the cell clusters; “Average Expression” is the z-score of the average gene expression of all cells within a cluster (scaled values); and “Percent Expressed” is the percentage of cells with non-zero gene expression.

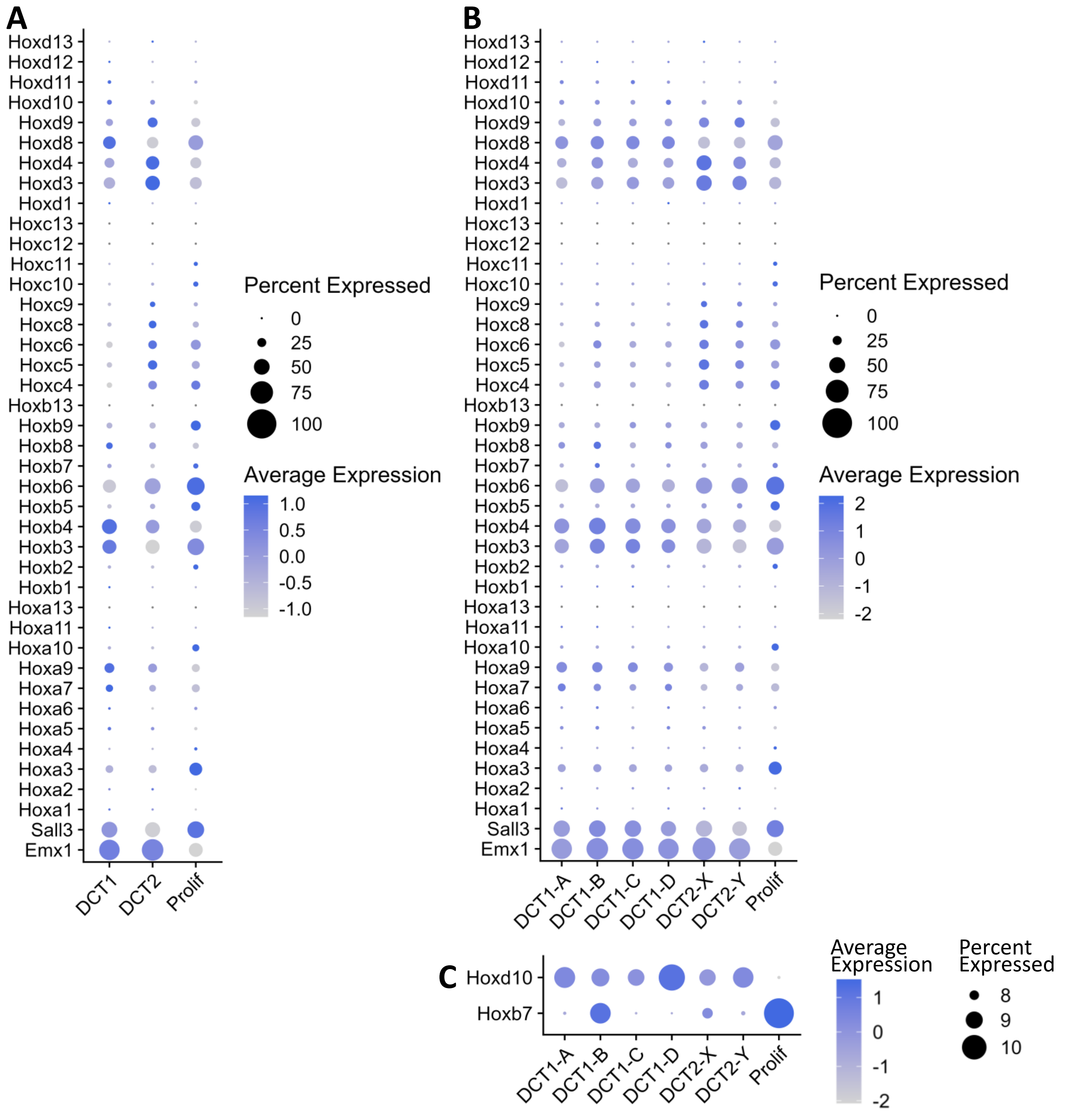


**Figure S14. Homeobox genes distribution.** Distribution of 39 homeobox transcripts, *Sall3* and *Emx1* in A) DCT1, DCT2, Proliferating cells, and in B-C) DCT1A-D, DCT2X-Y, and proliferating cells. Data are normalized and scaled (z-score) to examine relative expression across the cell clusters; "Average Expression" is the z-score of the average gene expression of all cells within a cluster (scaled values); and "Percent Expressed" is the percentage of cells with non-zero gene expression.
