## Supplemental Table 1 for "Enriched Single-Nucleus RNA-Sequencing reveals unique attributes of distal convoluted tubule cells"

Supplemental table 1. Key resources.

| REAGENT or RESOURCE | SOURCE | IDENTIFIER |
|---|---|---|
| **Antibodies** | | |
| Ki67, rabbit | Abcam | Cat#ab16667 |
| tNCC, rabbit | D.H.E. laboratory | |
| pNCC, rabbit | D.H.E. laboratory | |
| Calbindin D28k, mouse | Swant | Cat#CB300 |
| Parvalbumin, guinea pig | Swant | Cat#GP72 |
| GFP-FITC conjugated | Rockland | 600-102-215 |
| **Chemicals** | | |
| 1x DPBS | Gibco | Cat#14190-144 |
| 1x PBS | Gibco | Cat#20012-027 |
| cOmplete ULTRA Tablets, Mini, EDTA-free | Roche | Cat#5892970001 |
| RNasin Plus Ribonuclease Inhibitors | Promega | Cat#N2615 |
| SUPERaseIN RNase Inhibitor | Thermo Fisher Scientific | Cat#AM2696 |
| Protector RNase inhibitor | Sigma | Cat#PN 3335399001 |
| Vybrant™ DyeCycle™ Ruby Stain | Thermo Fisher Scientific | Cat#V10309 |
| Trypan Blue | Invitrogen | Cat#T10282 |
| Tissue-Plus™ O.C.T. Compound | Thermo Fisher Scientific | Cat#23-730-571 |
| ProLong® Diamond Antifade Mountant | Thermo Fisher Scientific | Cat#P36970 |
| Bovine serum albumin (BSA) | Thermo Fisher Scientific | Cat#BP1600 |
| DAPI | Sigma-Aldrich | Cat#D9542 |
| Tris-HCl | Thermo Fisher Scientific | Cat#BP153 |
| Collagenase, Type II | Thermo Fisher Scientific | Cat# 17101-015 |
| Gibco™ Leibovitz's L-15 Medium | Thermo Fisher Scientific | Cat#11-415-064 |
| Poly-D-lysine hydrobromide | Sigma | Cat#P7405 |
| **Critical commercial assays** | | |

| | | |
|---|---|---|
| Chromium Single Cell 3' GEM, Library & Gel Bead Kit v3 | 10x Genomics | Cat#PN-1000075 |
| Chromium Single Cell B Chip Kit | 10x Genomics | Cat#PN-1000153 |
| **Other supplies** | | |
| KONTES Dounce Tissue Grinders | Kimble Chase | Cat#KT885300-0002 |
| pluriStrainer 200 µm | pluriSelect | Cat#43-50200 |
| pluriStrainer 40 µm | pluriSelect | Cat#43-50040 |
| pluriStrainer 5 µm | pluriSelect | Cat#43-50005 |
| Connector Ring | pluriSelect | Cat#41-50000 |
| Falcon polystyrene Round-bottom Tubes 5mL | Thermo Fisher Scientific | Cat#352063 |
| Falcon Polystyrene Conical Tube 15 mL | Thermo Fisher Scientific | Cat#339650 |
| Falcon Polystyrene Conical Tube 50 mL | Thermo Fisher Scientific | Cat#339652 |
| Fuchs-Rosenthal disposable hemocytometer | INCYTO | Cat#DHC-F015 |
| DNA LoBind Tubes 1.5 ml | Eppendorf | Cat#022431021 |
| **Deposited data** | | |
| Raw and integrated snRNA-seq data | GEO | |
| Code | GitHub | |
| **Experimental models: Organisms/strains** | | |
| C57BL/6J | The Jackson Lab | Strain# 000664 |
| Slc12a3-IRES-CRE-ERT2 | The Jackson Lab | Strain# 030602 |
| CAG-Sun1/sfGFP | The Jackson Lab | Strain# 021039 |
| **Software and algorithms** | | |
| Cell Ranger | 10x Genomics | https://github.com/10XGenomics/cellranger |
| Seurat V4.0 | Satija Lab | https://satijalab.org/seurat/ |
| RStudio | RStudio, PBC | https://www.rstudio.com/ |
| IMARIS | Oxford instruments | https://imaris.oxinst.com/ |
| GraphPad Prism | Dotmatics | https://www.graphpad.com |
| ZEN | Zeiss | https://www.zeiss.com/microscopy/en/products/software/zeiss-zen-desk.html |
| Fiji ImageJ | NIH | https://imagej.net/software/fiji/ |
| **Mouse diets** | | |
| Control diet | Envigo | Cat#TD.190005 |
