## Supplemental Table 2 for "Enriched Single-Nucleus RNA-Sequencing reveals unique attributes of distal convoluted tubule cells"

| | | NK1 | NK2 | NK3 |
|---|---|---|---|---|
| Sequencing | Number of Reads | 180,789,479 | 355,942,388 | 379,966,834 |
| | Number of Short Reads Skipped | 0 | 0 | 0 |
| | Valid Barcodes | 95% | 93.50% | 93.60% |
| | Valid UMIs | 99.80% | 99.90% | 99.90% |
| | Sequencing Saturation | 45.70% | 54.00% | 54.10% |
| | Q30 Bases in Barcode | 94.80% | 96.20% | 96.20% |
| | Q30 Bases in RNA Read | 90.60% | 90.00% | 90.30% |
| | Q30 Bases in UMI | 94.40% | 95.70% | 95.60% |
| Mapping | Reads Mapped to Genome | 86.80% | 81.00% | 84.10% |
| | Reads Mapped Confidently to Genome | 84.30% | 77.50% | 81.00% |
| | Reads Mapped Confidently to Intergenic Regions | 4.60% | 5.40% | 5.50% |
| | Reads Mapped Confidently to Intronic Regions | 48.40% | 46.70% | 49.20% |
| | Reads Mapped Confidently to Exonic Regions | 31.30% | 25.40% | 26.30% |
| | Reads Mapped Confidently to Transcriptome | 51.10% | 44.20% | 45.80% |
| | Reads Mapped Antisense to Gene | 28.30% | 27.70% | 29.40% |
| Cells | Estimated Number of Cells | 10,189 | 14,003 | 13,346 |
| | Fraction Reads in Cells | 91.20% | 93.60% | 93.80% |
| | Mean Reads per Cell | 17,744 | 25,419 | 28,470 |
| | Median Genes per Cell | 1,652 | 1,862 | 2,014 |
| | Total Genes Detected | 24,048 | 24,870 | 25,351 |
| | Median UMI Counts per Cell | 3,393 | 3,948 | 4,391 |
| Sample | Chemistry | Single Cell 3' v3 | | |
| | Include introns | TRUE | | |
| | Reference Path | …refdata-gex-mm10-2020-A | | |
| | Transcriptome | mm10-2020-A | | |
| | Pipeline Version | cellranger-6.0.2 | | |
